# Deep Transfer Learning for Dormancy and Outbreaking State Classification in Metastatic Breast Tumor Cells: A Benchmark of Modern Deep Learning Models

**DOI:** 10.64898/2026.06.22.733720

**Authors:** Omveer Sharma, Keren Weidenfeld, Dalit Barkan, Oren Gal

## Abstract

Breast cancer cells that disseminate to distant organs can remain dormant (non-proliferative) for years before reactivating and progressing into lethal metastatic disease. Understanding the transition between dormancy and reactivation is therefore critical for early intervention and treatment. In this study, we investigate a comprehensive range of deep learning (DL) architectures to classify dormant versus proliferative breast tumor cells within a 3-dimensional growth factor reduced basement membrane extract (3D BME) system that models tumor dormancy and outgrowth. To capture the underlying spatiotemporal dynamics, we evaluate both spatial and sequence-based learning approaches. We consider convolutional neural networks (EfficientNet, ResNet, DenseNet, MobileNet, VGG, AlexNet), segmentation-based models (U-Net, U-Net++, Attention U-Net, DeepLabV3, HRNet) and transformer-based architectures (Vision Transformer, Swin Transformer, SegFormer). We investigate transfer learning using both fixed and fine-tuned strategies. Experimental results show that classification performance is greatly enhanced through the integration of temporal information. EfficientNet-B7, EfficientNet-B6, DenseNet-169, and DenseNet201 are consistently better than competing architectures for all tested models. EfficientNet-B7 with the use of temporal sequences input reaches an accuracy of 98.86% with a ROC-AUC of 0.998. The results highlight the significance of spatio-temporal feature learning and the value of DL frameworks in automated classification of dormant versus proliferative breast cancer cells in physiologically relevant microenvironments.

## I. Introduction

Breast cancer remains a global health challenge, affecting millions of women worldwide because of its highly complex and heterogeneous nature. The disease develops when abnormal breast cells proliferate uncontrollably and form tumors.

Given the molecular heterogeneity of breast cancer, early diagnosis, accurate subtype characterization, and monitoring of disease progression are essential for guiding personalized treatment decisions and optimizing clinical outcomes and survival [1]. Despite significant advances in diagnosis and treatment, metastasis (the spread of tumor cells to other organs in the body) remains the leading cause of breast cancer-related mortality [2–5]. Metastatic disease may be detected at diagnosis or can emerge years or even decades after the primary tumor has been removed and initial treatments have been completed. The long latency period of recurrent metastatic disease is clinically attributed to the presence of dormant (quiescent) disseminated tumor cells. These cells, which spread early from the primary tumor and colonize specific distant organs, remain dormant and asymptomatic until they are reactivated and develop into overt metastases [6]. Dormant tumor cells are strongly associated with therapy resistance, metastatic relapse and poor long-term patient outcomes [7–9]. These hibernating cells escape immune-surveillance and launch adaptive stress-response mechanisms [4]. This allows dormant cells to survive for years or even decades in distant tissues such as bone marrow, lungs, liver and brain. Tumor microenvironment plays important roles in dormancy and reawakening of metastasis [9]. Stromal and immune cells are key components of the metastatic niche that support dormant disseminated tumor cells (DTCs) and regulate their reactivation. Together with tumor cells, these microenvironmental constituents actively remodel the extracellular matrix (ECM), altering its composition, organization, and mechanical properties. Such ECM remodeling creates a dynamic microenvironment that can either maintain DTCs dormancy or promote their awakening and subsequent metastatic outgrowth [10].

To investigate the role of the microenvironment in regulating tumor cell dormancy and metastatic progression, Barkan and colleagues developed previously a three-dimensional (3D) reconstituted basement membrane extract (BME) system that models dormancy and metastatic outgrowth in vitro under physiologically relevant microenvironmental condition [11, 12]. BME represents a specialized form of ECM that recapitulates key structural and biochemical features of the basement membrane. This model has provided important insights into how ECM and microenvironment derived cues govern the balance between tumor cell quiescence and proliferation [13]. A notable advantage of this system is that it enables realtime monitoring of tumor cell dormancy and metastatic outgrowth through time-lapse microscopy, allowing the dynamic behavior of individual cells to be tracked in well-controlled microenvironmental settings. The resulting image datasets provide a powerful platform for quantitative analysis of cellular state transitions. Deep learning (DL) has emerged as a highly successful approach in biomedical image analysis, enabling the automatic learning of complex feature representations directly from imaging data [14]. Convolutional neural networks (CNNs) such as ResNet and EfficientNet have shown great performance in large scale image classification and medical imaging applications [15–19]. In a similar vein, segmentation frameworks such as U-Net, DeepLab and HRNet have become standard solutions to pixel-level biomedical image analysis tasks [20–22]. More recently, transformer-based architectures have exhibited a remarkable ability to model global contextual relationships and long-range dependencies in computer vision tasks. Vision Transformer (ViT), Swin Transformer and SegFormer use the attention mechanism to learn hierarchical representations, and they have achieved state-of-the-art performance on classification and segmentation applications [23–25]. The emergence of large-scale vision models like Segment Anything and ConvNeXt V2 [26, 27] emphasizes the importance of scalable and transferable visual representations in biomedical imaging.

However, despite these advances, most of the existing DL studies in cancer imaging are focused mainly on static image analysis and ignore the intrinsic temporal dynamics of cancer-cell behavior. Biological processes like dormancy, activation, migration and proliferation evolve continuously in time and therefore require models that can learn long range temporal dependencies. Although temporal modeling has brought great benefits to video understanding and medical time-series analysis, its applications to cancer dormancy analysis are still limited [8, 28, 29]. Existing approaches often miss subtle spatiotemporal transitions associated with dormant-to-proliferative state changes, particularly in physiologically relevant 3D microenvironmental systems. This limitation limits the ability of current computational methods to fully explain the dynamic evolution of metastatic cancer cells.

In this work we address these issues by systematically exploring DL architectures for the classification of dormant and proliferative breast cancer cells in a 3D BME system. We explore both spatial and temporal modeling strategies to capture the underlying spatiotemporal dynamics of cancercell behavior across image sequences. Specifically, we systematically evaluate convolutional, segmentation-based, and transformer-based architectures under consistent experimental settings while also analyzing the impact of transfer learning strategies. Our experiments show a significant improvement in the classification performance by incorporating temporal information, highlighting the importance of modeling the dynamic cellular behavior. Among the evaluated models, EfficientNet-based architectures consistently achieve the best performance, with EfficientNet-B7 obtaining an accuracy 98.86% and a ROC-AUC of 0.998. These results demonstrate the power of spatiotemporal DL approaches for the automation of cancer dormancy analysis and provide a basis for future investigation of metastatic cancer-cell dynamics in physiologically relevant environments with artificial intelligence.

## II. Related Work

In this section, we review recent advancements in breast cancer image classification, where data-driven approaches have been adopted. Broadly, these methodologies can be categorized as machine learning (ML) based approaches and deep learning (DL) based approaches.

### A. Classification using ML Approaches

Classification of histological images of breast cancer is an important element of research in the field of medical imaging. For high-precision algorithms such as K-Nearest Neighbors (K-NN), Random Forest (RF), Naive Bayes (NB) and Support Vector Machines (SVM) have been used for this task. Algorithms extract features such as shape, texture and density from histological images and classify them as malignant or benign. In the case of high dimensional data and complex boundary of the classes, SVMs are preferable. The RF combine several projections of decision trees for better robustness. K-NN classifies data based on spatial proximity, whereas NB classifies data based on probabilistic relationships of features and class labels [30].

### B. Classification using DL Approaches

The DL approaches have been predominantly adopted. Unlike traditional ML, which relies on manual feature engineering, DL models automatically extract discriminative features from complex medical images [31, 32]. Recent advances in breast cancer analysis can generally be categorized into CNN-based methods, transformer-based architectures, and hybrid frameworks that combine convolutional and attention mechanisms. CNN-based methods are very good at capturing local spatial patterns and texture information, while transformer-based models are better at modeling long-range contextual relationships due to attention mechanisms. More recently, hybrid architectures of CNNs and transformers were proposed to achieve better performance by exploiting the advantages of both local feature extraction and global contextual learning.

#### 1) ResNet

Researchers proposed the Multi-Strategy Parrot Optimizer (MSPO) combined with ResNet18 to enhance classification accuracy on the BreaKHis breast cancer image dataset [33]. ResoMergeNet (RMN), a proposed multi-modal classification framework, enhances diagnostic accuracy for conditions such as cataracts and cancer. RMN employs transfer learning (pre-trained ResNet-50) and specialized ResBoost and ConvMergeNet modules to efficiently extract and fuse features from different sources [34]. Zhang et al. developed a ResNet-based DL framework that combined full-breast mammograms with preoperative clinical variables to predict lymph node metastasis in early-stage breast cancer, and showed that the integrated model outperformed the models based on clinical variables only, showing a promising potential in reducing the number of unnecessary sentinel lymph node biopsies [35].

#### 2) Transformer

In recent years, transformer-based models have been widely used for medical image classification tasks due to their powerful attention mechanism, which demonstrates strong performance and improved ability to capture complex spatial relationships in medical data [36]. Malik et al. used a multi-modal dataset that included clinical, molecular, demographic, and lifestyle information, and proposed a transformer-based survival module and an attention-guided classification module for identifying second primary cancer, low-risk recurrence, and high-risk recurrence [37]. An anatomy-aware multiview transformer framework is proposed for breast microcalcification classification using dualview mammographic images while preserving fine-grained microcalcification texture patterns and imposing anatomical consistency between CC and MLO views, with high diagnostic performance on the CBIS-DDSM dataset while improving the interpretability with clinically relevant visual explanations [38].

#### 3) Hybrid model

A hybrid model, namely CTNet, is proposed that integrates CNN with ViT to enhance breast cancer classification. It consists of a channel attention module for refining modality-specific features, a Cross Attention Block (CAB) for efficient feature extraction, and a Dynamic Attention (DA) block combined with Transformer encoding for improving contextual representation [39]. Israa et al. proposed a hybrid DL framework for classifying multi-class BI-RADS breast cancer from mammographic images [40]. This dual-stream architecture integrates the lightweight CNN MobileNetV1 to extract the fine-grained local features and the ViT to learn global contextual relationships, which enables the complementary representation learning. The study proposed TransBreastNet, a hybrid CNN-Transformer DL framework for the joint breast cancer subtype classification and temporal lesion progression analysis from mammogram images [41]. This framework combines spatial lesion features, temporal progression patterns and patient-specific clinical metadata in a multimodal multitask architecture, and achieves high performance in subtype and stage prediction with greater interpretability and clinical relevance. A Transformer-based Hybrid framework was proposed for automatic detection of breast cancer and its subtype identification from whole slide pathological images [42]. This framework integrates CNN-based models, ViT, and voting, and achieved the good performance for both cancer detection and subtype classification, and improved the diagnostic accuracy and reduced the workload of pathologists.

## III. PRELIMINARIES

In this section, we discuss the key concepts and mathematical background required for the proposed framework. First, the problem formulation is presented to define the goal of classifying dormant versus proliferative cancer cells from temporal image sequences. Then, we introduce the basic principles of CNNs to learn long-range temporal dependencies in sequential biomedical data.

### A. Data Preparation and Problem Formulation

For this work images were taken at 30 minute intervals for about 145 timestamps per sequence. D2OR/D2A1 cells (2000 cell/well) were seeded in 96 well plate, pre-coated with 50µl of Cultrex growth factor-reduced basement membrane extract (BME, Trevigen) per well. After seeding the plates were transferred to Incucyte® SX5 device (Sartorius) for automated, incubator based live cell analysis. Scanning was performed using “Organoid assay” scan type, in 4X objective, one image per well in 30 min intervals for 72 hours. Brightfield images were exported as series of Tiff images and aligned using NIS element Ar software Ver.4.20. The fullsize images of 1040*×*1408 pixels were manually cropped into two different sizes of sub-images of 96*×*96 and 64*×*64 pixels. For the first timestamp, the target cell was centered in the sub-image. We selected these two specific dimensions to assess the best bounding region that can continuously contain the cells from the first timestamp to the last timestamp. A larger dimension of 128*×* 128 was omitted to avoid adding an unnecessary computational burden. During the manual cropping process, we observed that with the 64*×* 64 resolution in the manual cropping process, cells grew to the image edges over time in some samples and 96*×* 96 dimension was more appropriate. However, the DL models were tested rigorously on both dimensions to empirically investigate the impact of insufficient spatial coverage on classification performance as cells progressively outgrow their localized regions.

The whole dataset has two classes of cancer-cell trajectories, dormant cells and outbreaking cells. Experiments were carried out to evaluate the effect of the image resolution and the temporal context on the classification performance considering the image sizes of 64*×*64 and 96 *×* 96 pixels. For each temporal configuration, the data were independently partitioned into training, validation, and testing subsets. The full-length sequence setting (*≈*145) 64*×*64 dataset includes 3331 samples, of which 1807 are dormant-cell samples and 1524 are outbreaking-cell recordings. These samples were divided into 2331 training, 500 validation and 500 testing samples, as detailed in Table I. The 96 *×*96 dataset has 3787 samples, consisting of 2058 dormant-cell samples and 1729 outbreaking-cell samples. The training, validation and testing set were allocated with 2651, 568 and 568 samples, respectively.

**TABLE I:**
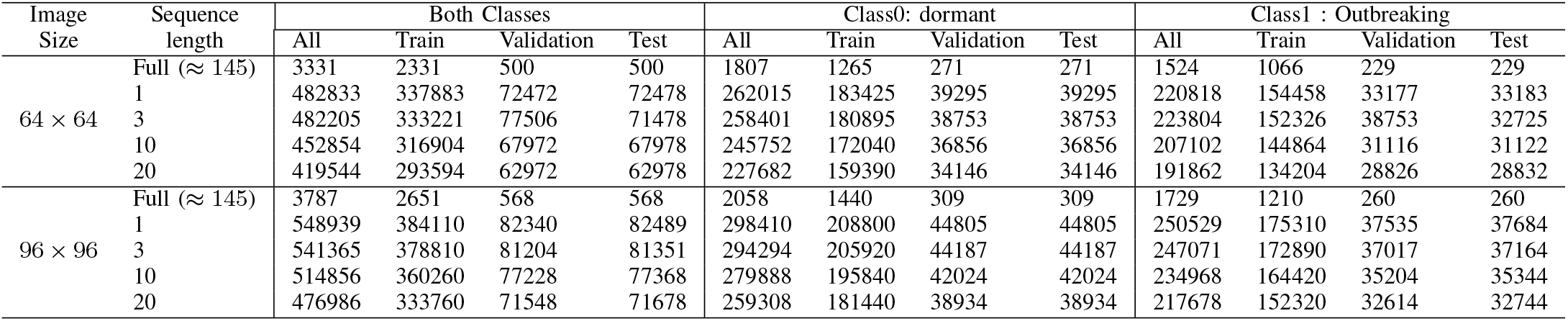
Dataset distribution for different temporal sequence configurations, including the total number of samples and the corresponding training, validation, and testing splits for dormant and outbreaking cancer-cell classes.

To investigate the impact of temporal information, the original long-term recordings were segmented into shorter sequences of lengths 1, 3, 10, and 20 timestamps. For the 64 *×*64 image resolution, the resulting datasets contain 482833, 482205, 452854, and 419544 samples for sequence lengths of 1, 3, 10, and 20, respectively. For the 96*×* 96 image resolution, the corresponding datasets contain 548939, 541365, 514856, and 476986 samples. For each temporal configuration, the data were split into training, validation, and testing subsets using an approximate ratio of 70%, 15%, and 15%, respectively. To maintain class balance, dormant and outbreaking samples were partitioned separately using the same ratio. Table I gives a detailed description of the training, validation and testing samples per class, image resolution and temporal configuration.

Let the input dataset be represented as:

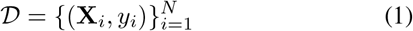

where *N* is the total number of samples, **X**_*i*_ denotes the temporal image sequence of the *i*^*th*^ sample, and *y*_*i*_ *∈ {*0, 1} is the class label, where 0 and 1 correspond to dormant and proliferative states respectively. Each temporal sample consists of a sequence of microscopy images:

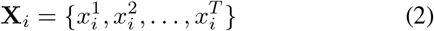

where *T* is the number of temporal frames and each frame 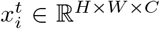 represents a grayscale image with height *H*, width *W*, and *C* = 1 channel.

The objective is to learn a mapping function:

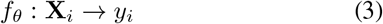

where *θ* are the learnable parameters of the model. The model should be able to capture spatial cellular morphology and temporal progression dynamics after training to accurately identify dormant and proliferative cancer-cell behaviors.

The representative samples shown in Fig. 1 illustrate the visual appearance of both classes at the beginning and end of the temporal sequence. The comparison of the first frame (*t* = 1) and the last frame (*t* = *T≈* 145) reveals the temporal variations and structural variations over the sequence. It should be noted that it is relatively difficult to differentiate between the two classes from the first frame, since both classes have highly similar visual properties. On the other hand, the last frames show more obvious and observable differences between the classes. The observation indicates that the DL models may have difficulty in correctly differentiating the classes with the early temporal information only and benefit from learning the temporal dynamics over the whole sequence.

**Fig. 1:**
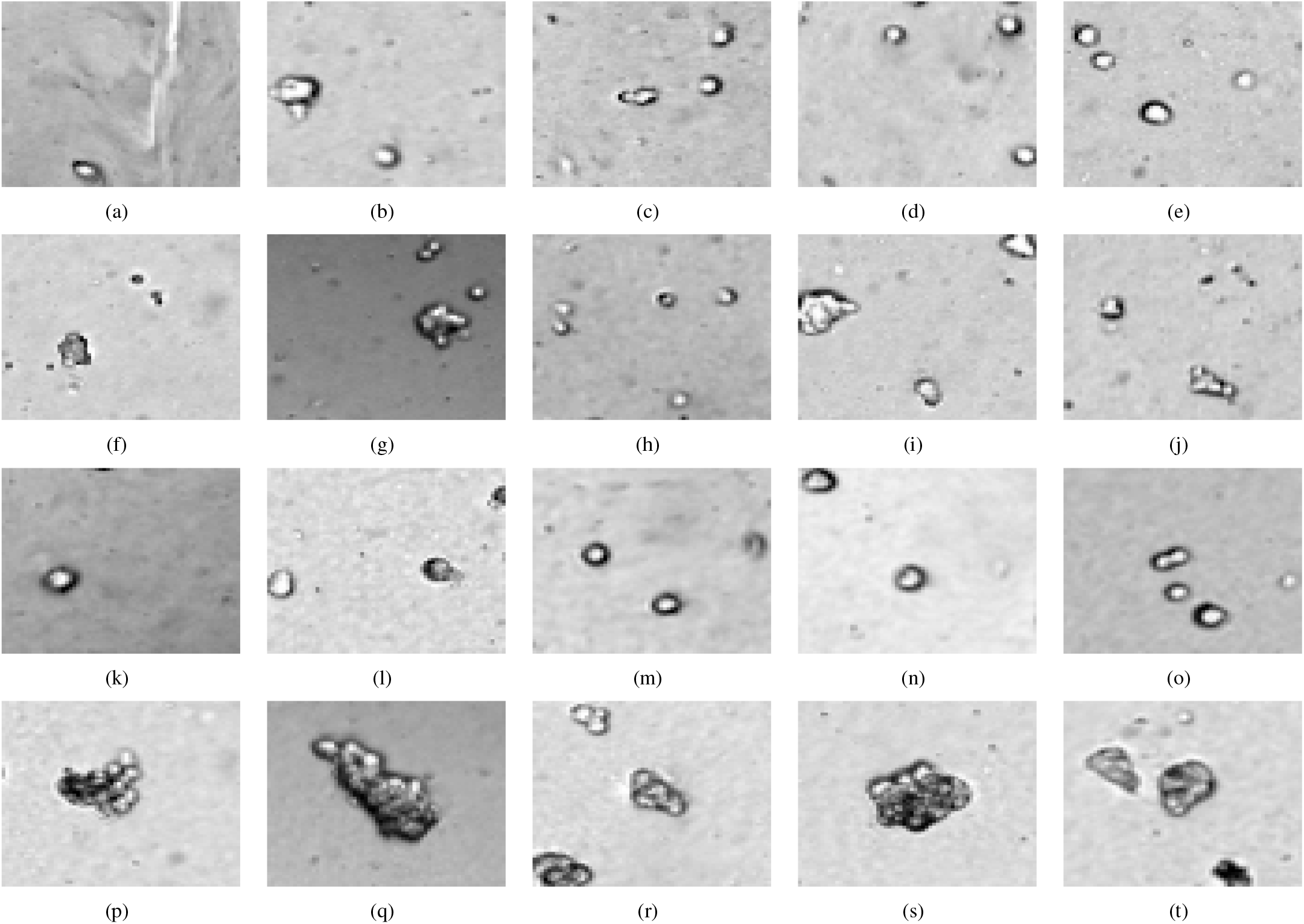
Representative examples from the dataset showing temporal changes between the beginning and end of the sequence for both classes. Figures 1(a)–1(e) illustrate the first temporal frame (*t* = 1) for samples belonging to class 0, while Figures 1(f)–1(j) show the first temporal frame (*t* = 1) for samples from class 1. Similarly, Figures 1(k)–1(o) present the final temporal frame (*t* = *T*, where *T≈* 145) for class 0 samples, whereas Figures 1(p)–1(t) correspond to the final temporal frame (*t* = *T*, where *T≈* 145) for class 1 samples. These examples highlight the visual and temporal variations between the two classes across the sequence progression.

### B. Convolutional Neural Networks (CNNs)

The CNNs are among the most popular DL architectures for image analysis and can learn hierarchical spatial representations from the raw image data automatically. CNNs operate by training trainable convolutional filters to slide along the spatial dimensions of an input image, learning local patterns such as edges, textures, shapes, and cellular morphological structures. CNNs learn feature representations that are more abstract and discriminative after multiple convolutional layers.

For an input feature map **X** and convolution kernel **K**, the convolution operation can be expressed as:

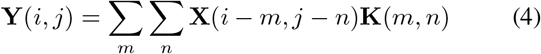

where **Y**(*i, j*) is the output feature value at the spatial location (*i, j*), **X** is the input feature map and **K** is the learnable convolution kernel. The convolution operation enables the network to extract information from a local neighborhood as well as to preserve spatial relations in the image.

A typical CNN architecture is composed of convolutional layers, nonlinear activations, pooling operations, normalization layers and fully connected layers. Pooling operations reduce spatial dimensionality while retaining salient semantic information, leading to improved computational efficiency and robustness to spatial variations. CNNs have demonstrated a strong capacity in biomedical imaging applications to extract fine-grained spatial features corresponding to cellular morphology, tissue organization, and cancer-related structural patterns [15–17].

### C. Multi-Head Self-Attention (MHSA)

Recently, transformer-based architectures have achieved great success on computer vision tasks thanks to their ability to capture long-range spatial dependencies and global contextual information. Transformer models use self-attention mechanisms to relate all the input elements to each other at the same time. Multi-Head Self-Attention (MHSA) introduced in the Transformer architecture by Vaswani et al. [43] allows the network to learn multiple feature representations in parallel and improve the contextual understanding.

Given an input feature matrix: **X**∈ ℝ^*N ×d*^. Where *N* is the number of input tokens and *d* is the feature dimension, three feature projections are produced:

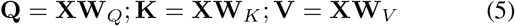

where **Q, K**, and **V** are the query, key, and value matrices respectively and **W**_*Q*_, **W**_*K*_, and **W**_*V*_ are trainable projection matrices.

The scaled dot-product attention operation is defined as:

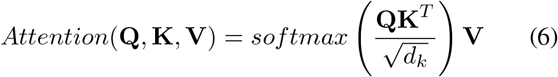

Here *d*_*k*_ is the dimension of key vectors and the term 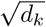 is a scaling factor that prevents the values from getting too large during optimization.

MHSA breaks the feature space into multiple attention heads instead of learning a single attention representation:

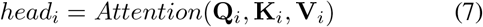

where *i* = 1, 2, …, *h* and *h* is the total number of attention heads.

The outputs of all the attention heads are concatenated and projected to get the final representation:

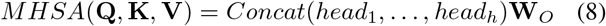

where **W**_*O*_ is the learnable output projection matrix.

The multiple attention heads enable transformer models to attend to different spatial relations and feature properties at the same time. This property allows architectures like ViT, Swin Transformer and SegFormer to effectively capture local and global context dependencies. MHSA has shown promising ability of learning complex structural patterns and improving feature representation for classification and segmentation tasks in biomedical imaging applications [23–25].

## IV. Network Architecture

In this section, we outline the baseline architectures considered in this work and the transfer-learning configurations adopted for model adaptation. In a systematic attempt to study the efficacy of different DL paradigms to classify dormant versus proliferative breast cancer cells, we evaluate CNN- based models, segmentation-based architectures, and transformer-based networks. To compare fairly, all models are trained on the same experimental conditions.

### A. Baseline Models

To comprehensively evaluate spatial and spatio-temporal representation learning, DL architectures of multiple categories were taken into consideration. The following Models are incorporated in this study.

- **ResNet-18, ResNet-34, ResNet-50, and ResNet-101** These models are constructed on residual learning and CNN based classification architectures [15]. ResNet employs skip connections, which help to solve the vanishing gradient problem and allow very deep networks to learn more discriminative hierarchical feature representations. The residual structure can help to optimize efficiently and propagate features to multiple layers. These advantages have led to the success of ResNet based models in many computer vision and biomedical imaging tasks. To enhance the predictive performance and clinical utility, ResNet-based frameworks for breast lesion classification and medical image diagnosis have been utilised [35].
- **DenseNet-121, DenseNet-169, and DenseNet-201**

The models are based on densely connected CNNs [44]. DenseNet connects each layer to every other layer in a feed-forward way, thus improving the flow of information and the gradient throughout the network. Such dense connectivity reduces the number of trainable parameters while preserving strong representational power. The architecture has shown remarkable effectiveness in medical imaging applications. DenseNet is employed in the CheXNet framework to perform automatic detection of chest diseases based on radiographic images, and reported performance on par with radiologists [17].

- **EfficientNet-B0, EfficientNet-B2, EfficientNet-B3, EfficientNet-B4, EfficientNet-B5, EfficientNet-B6, and EfficientNet-B7**

These models utilize compound-scaled CNN architectures [16]. EfficientNet proposes a compound scaling method, which uniformly scales network depth, width, and resolution. This improves accuracy with the same computational complexity. The balanced scaling allows the network to learn rich hierarchical representations with less parameters than conventional CNN architectures. EfficientNet models have been shown to perform well on multiple biomedical image analysis tasks, such as breast cancer histopathological image classification [31].

- **MobileNet-V2, MobileNet-V3-Large, and MobileNetV3-Small**

Lightweight CNN architectures for computationally efficient image recognition tasks [45, 46]. The MobileNet architectures employ depth-wise separable convolutions and inverted residuals to greatly reduce the computational cost of the network while preserving the predictive performance. They are well suited for low resource settings and health applications, being efficiently designed. Previous work has shown the successful use of MobileNet-based DL frameworks for disease classification and medical image analysis tasks [32].

- **U-Net, U-Net++, Attention U-Net, Efficient Attention U-Net, DeepLabV3, HRNet, ResU-Net, and Cellpose** These models are mainly segmentation-based architectures for learning detailed spatial representations from biomedical images. U-Net proposed has an encoderdecoder framework with skip connections to preserve spatial information [20]. U-Net++ further improved feature fusion by adding nested dense skip pathways [47]. Attention U-Net utilised attention mechanisms to focus on relevant spatial regions [48]. DeepLabV3 extracted multi-scale features with atrous convolution [21]. HRNet preserved the high-resolution feature representation across all network layers [22]. Cellpose was developed for generic cell segmentation across a broad set of microscopy datasets [49]. These architectures have been used to achieve excellent performance in cell detection, tissue segmentation, tumour delineation, and other biomedical imaging tasks.
- **Vision Transformer (ViT), Swin Transformer, and SegFormer**

These are transformer architectures developed for learning visual representations using self-attention mechanisms. Dosovitskiy et al. propose ViT [23] which applies transformer operations directly on image patches to learn global contextual relationships. Swin Transformer introduced shifted window attention to enhance computational efficiency in hierarchical feature learning [24]. SegFormer uses transformer encoders and light-weight decoders for semantic segmentation [25]. Transformer-based methods have shown promising performance on a variety of medical imaging tasks by modeling long-range spatial dependencies and contextual information [36].

### B. Transfer Learning Configurations

To study the effect of parameter adaptation on classification performance, transfer learning was used. Since the microscopy datasets used in this work are relatively smaller than largescale natural image datasets, pre-trained ImageNet weights were utilized as initialization for all applicable architectures. Four transfer-learning configurations were investigated:

#### 1) Configuration 1: Full Fine-Tuning

All network parameters were initialized using pretrained weights and updated during training. The complete model was retrained using the target dataset:

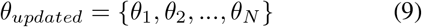

where all parameters remain trainable.

#### 2) Configuration 2: Training with Frozen Backbones

We froze the entire pre-trained backbone and only finetuned the first convolution layer adapting to the sequence and the final classification layer:

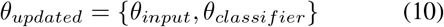

while all intermediate feature extraction layers were frozen.

#### 3) Configuration 3: Training on the Half-Model

The network was split into two parts, where only the first part of the model was extracted and retrained:

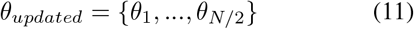

This allows for the adaptation of higher semantic level representations while preserving low-level learned features.

#### 4) Configuration 4: Partial Fine-Tuning

The model was divided into four equal parts, and only the two initial quarters were considered for adaptation. The first quarter was frozen while the remaining network layers were retrained:

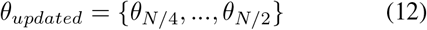

This configuration aims to balance feature preservation and model adaptability.

This configuration aims to balance feature preservation and model adaptability while reducing the number of trainable parameters. These transfer-learning configurations allow investigation of the influence of parameter freezing and feature adaptation on dormant versus proliferative breast cancer-cell classification.

### C. Training Configuration and Hyperparameters

The dataset was randomly split using a fixed random seed for all experiments into training (70%), validation (15%) and testing (15%) sets for a consistent and fair comparison among all the evaluated models. Input normalization was used during training while no data augmentation was performed. All the models were trained with Adam optimizer with the initial learning rate of 0.0005 and the categorical cross-entropy loss function. We trained with a batch size of 16 and a maximum of 50 epochs. The overfitting was reduced by applying a dropout rate of 0.1. The model selection is based on the validation performance where the best model is selected based on a validation metric which is a combination of overall accuracy and the class-wise accuracy. In order to enable efficient convergence, the learning rate was reduced by a factor of 0.5 if no improvement of validation performance was observed for 10 consecutive epochs. We also used early stopping, if the validation performance did not improve for 13 consecutive epochs, to avoid overfitting and unnecessary computation. The model parameters with best validation performance were saved and used for final testing and frame-wise evaluation. All models were trained and evaluated on an NVIDIA A100-SXM4-80GB GPU.

## V. Results and discussion

In this section, we analyze the performance of different DL architectures in classifying dormant and proliferative breast cancer cells from microscopy image sequences. The experiments investigate the effect of transfer-learning strategy, temporal sequence length and image resolution on classification performance. We tested both 64 *×*64 and 96*×*96 cropped image sizes in different temporal settings.

### A. *Transfer Learning Analysis on* 64 *×* 64 *Images*

The performance comparison of different DL architectures is shown in Table II for four transfer-learning configurations,namely C1, C2, C3 and C4 (as described in the previous section), using 64 *×* 64 image resolution. The table reports the overall classification accuracy together with its 99% confidence interval (CI99), class-wise performance in terms of recall (Class 1 accuracy) and specificity (Class 0 accuracy), and the corresponding ROC-AUC values. During these experiments, only one image is taken at a time; thus, the input sequence length is set to one. The experimental results show that the transfer-learning strategy has the ability to improve the classification performance.

**TABLE II:**
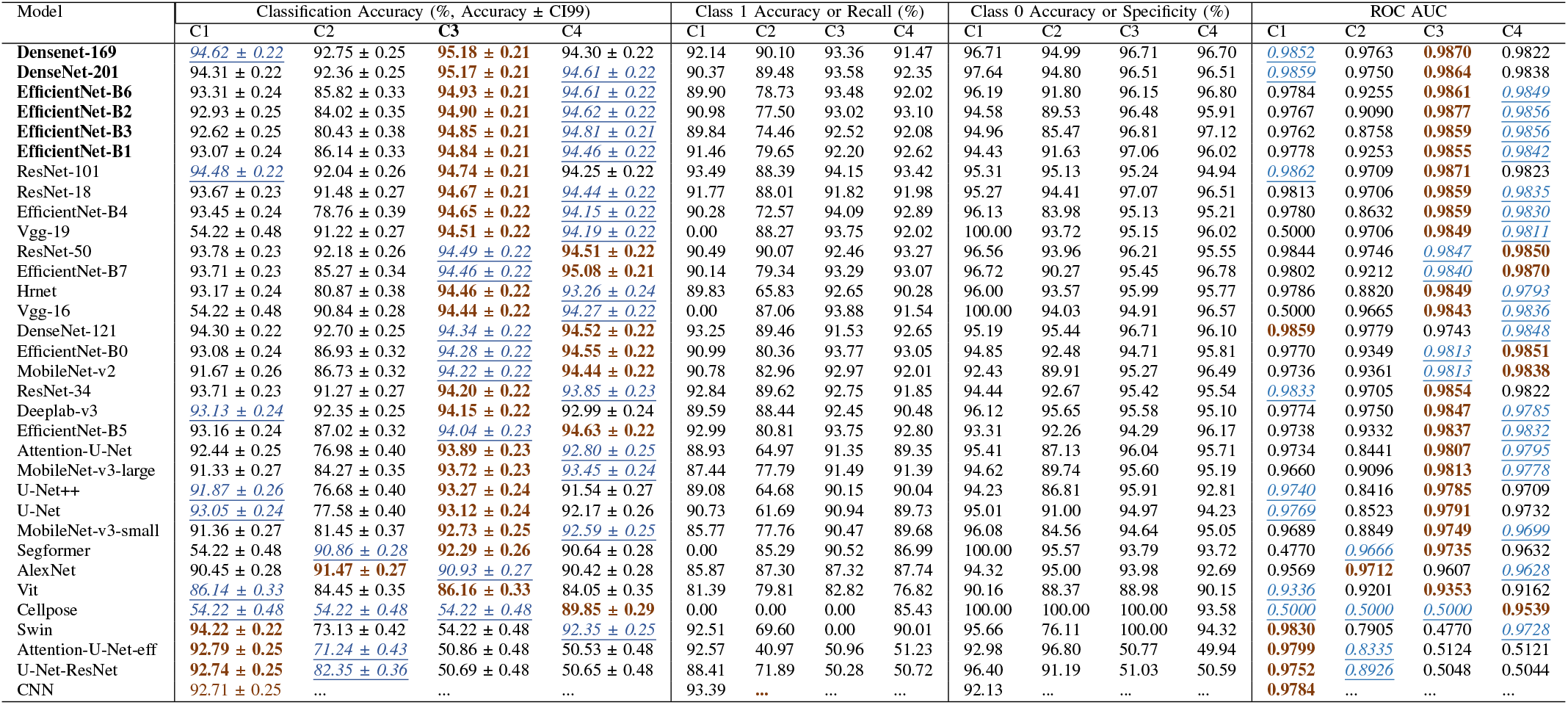
Performance comparison of advanced DL models based on classification accuracy and ROC metrics under configurations C1, C2, C3, and C4. For each model, the best-performing configuration in terms of accuracy is highlighted in **bold**, while the second-best configuration is indicated using *<u>italicized and underlined</u>* text.

Configuration C3 and C4 have performed better across most architectures amongst all evaluated configurations. In these settings, we re-trained the higher-level layers of the pre-trained networks with different label settings while keeping the lowlevel learned feature representations. This transfer learning strategy allowed the models to adapt well to the microscopy-specific cellular morphology, while preserving strong low-level spatial feature extraction. Specifically, C3 outperformed C4 in a more dominant and stable manner, as shown in Table II and Fig. 2. The Fig. 2 illustrates the overall classification accuracy comparison of configurations C1 to C4 across the top-performing 10 models. On the contrary, configuration C2 generally gave less favorable outcomes in most architectures. In this setting, most of the backbone was kept frozen and the models were not able to sufficiently adapt to the domain specific properties of microscopy images. C2 was particularly constrained in performance for EfficientNet and for architectures that employed segmentation. The C1 configuration did average to moderate on average across the board, as shown in Table II and Fig. 2.

**Fig. 2:**
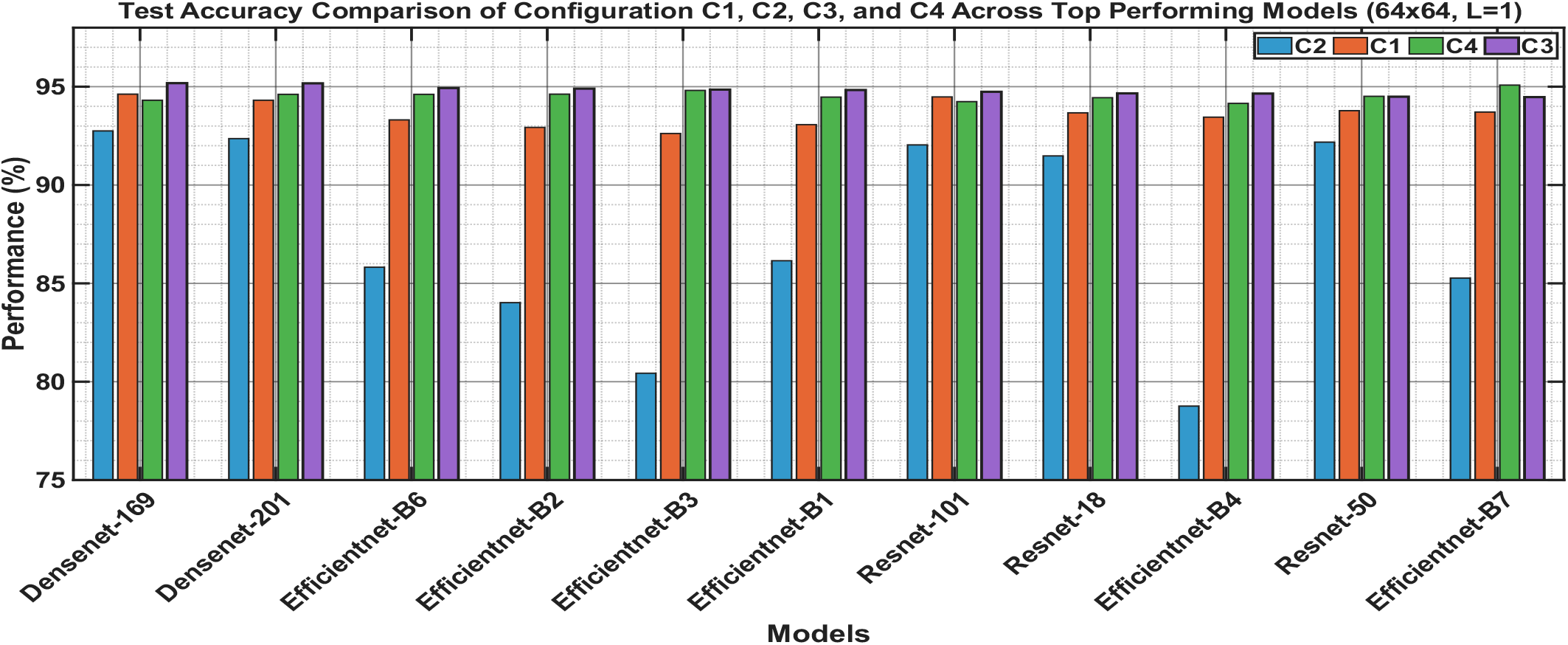
Comparison of Overall Accuracy Across Configurations C1–C4 for the Top Performing Models Using Single-Image Input (64 *×* 64).

The experimental results also indicate that EfficientNet and DenseNet based architectures always achieve better results than competing models for all transfer-learning settings. DenseNet-169 and DenseNet-201 achieved best results, with test accuracies 95.18 and 95.17%, respectively and ROC-AUC values close to 0.99, which shows the effectiveness of dense connectivity in feature propagation for biomedical image tasks. EfficientNet based model also achived good results.

In particular, EfficientNet-B6 achieved one of the highest performances with a test accuracy of 94.93% and ROC-AUC of 0.9861 on the Configuration C3. Similarly, EfficientNet-B2, EfficientNet-B3, EfficientNet-B4 and EfficientNet-B7 showed highly competitive performance, confirming the effectiveness of compound-scaled feature learning for the microscopy-based cancer-cell analysis.

Residual learning-based architectures such as ResNet-18, ResNet-34, ResNet-50 and ResNet-101 showed stable and competitive performance in all transfer learning settings. Un-der Configuration C3, ResNet-101 achieves the best performance among these with a test accuracy of 94.74% and ROC-AUC of 0.9871. These results demonstrate that residual feature learning can effectively capture discriminative cellular morphology and structural features. Additionally, segmentation-oriented architectures like U-Net, U-Net++, Attention U-Net, DeepLabV3 and HRNet showed good classification performance, despite being originally proposed for pixel-level segmentation tasks. DeepLabV3 and HRNet in particular achieved competitive performance due to their ability to preserve multiscale contextual information and high-resolution spatial representations. Also, attention-based segmentation architectures benefited from their ability to focus on biologically relevant cellular regions.

The performance on transformer-based architectures was relatively mixed. SegFormer achieved a moderate accuracy for classification, but the ViT performed worse than CNN-based architectures. In several hybrid transformer-based segmentation models, as well as on some configurations of the Swin Transformer, we observed unstable convergence behavior, which suggests that these architectures may require several times larger datasets for effective optimization. In summary, the results indicate that CNN-based architectures, in particular the EfficientNet and DenseNet families, offer the best balance between feature extraction capability, generalization performance and computational efficiency for dormant vs. proliferative cancer-cell classification. Overall, DenseNet-169, DenseNet-201, EfficientNet-B6, and EfficientNet-B2 are the best-performing models. The following section further analysis the input sequence-length effect over overall performance for configuration C3, as it performed better than the other architectures in most cases while sequence length is considered one.

### B. *Effect of Temporal Sequence Length on* 64 *×* 64 *Images*

In this section, we evaluate the prediction performance of all baseline models by sequentially increasing the input sequence length from 1 to 20. Specifically, sequence lengths of 1, 3, 10 and 20 were examined for all training configurations (C1, C2, C3 and C4). However, results for Configuration C3 are presented in this section, because C3 outperformed the other configurations in the previous experiments with the input sequence length set to one in consistent manner. For completeness, the results for the other three transferlearning configurations (C1, C2, and C4) are included in the supplementary material (Tables S1, S2 and S3).

Table III shows the effect of the temporal sequence length on the classification performance for the 64*×* 64 image resolution in configuration C3. The table reports the overall classification accuracy together with its 99% confidence interval, class-wise performance in terms of recall (Class 1 accuracy) and specificity (Class 0 accuracy), and the corresponding ROC-AUC values. The experimental results demonstrate that the models can greatly improve their ability to discriminate between proliferative and dormant cancer cell behaviours by including the temporal information. The Fig. 3 compares the best performing models for different temporal sequence lengths, which emphasizes the importance of temporal information for classification accuracy. We observed consistent improvements in classification accuracy and ROC-AUC as the sequence length increased from 1 to 3, then to 20 frames, for most architectures.

**TABLE III:**
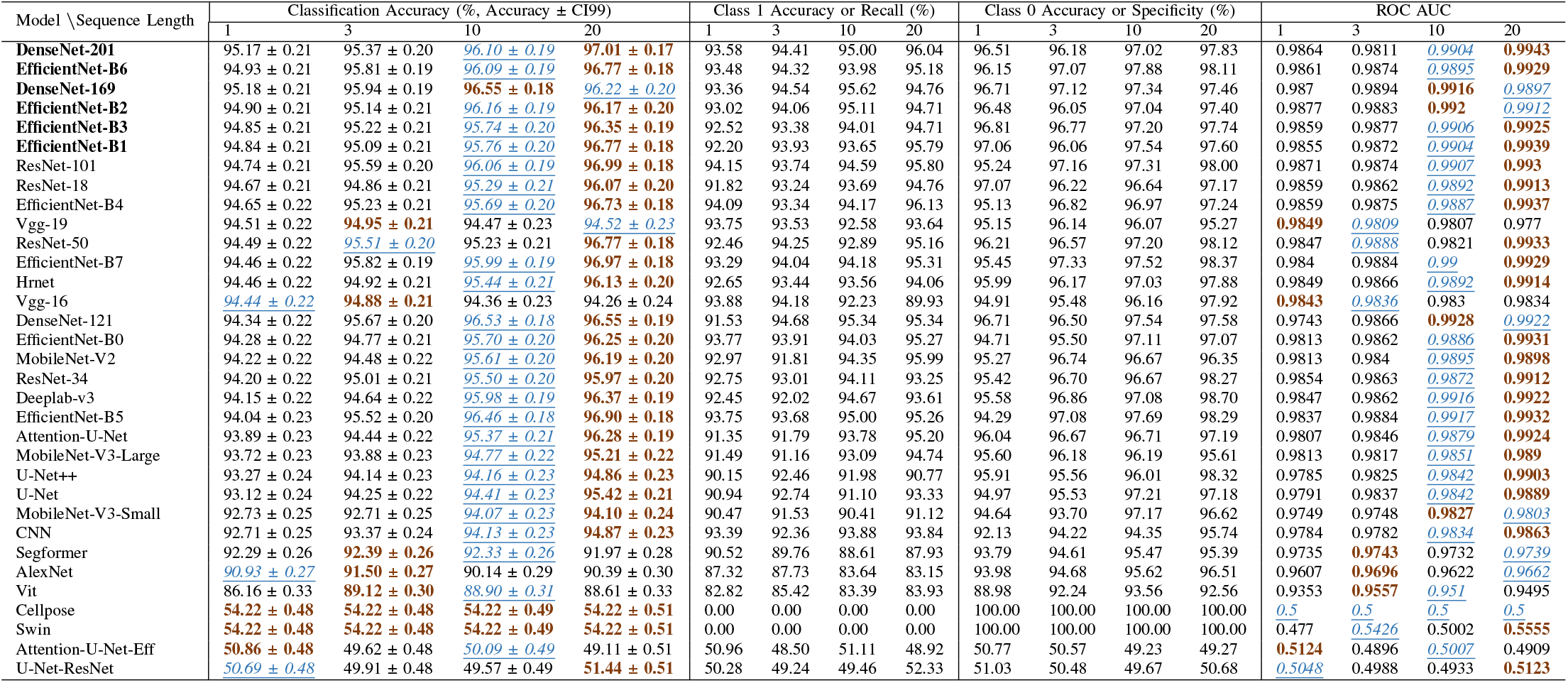
Performance Comparison of Advanced DL Models Based on Classification Accuracy and ROC Metrics Across Sequence Lengths 1, 3, 10, and 20 for 64*×* 64 Input Images. The best and second-best configurations for each model are highlighted in **bold** and *<u>italicized-underlined</u>* text, respectively.

**Fig. 3:**
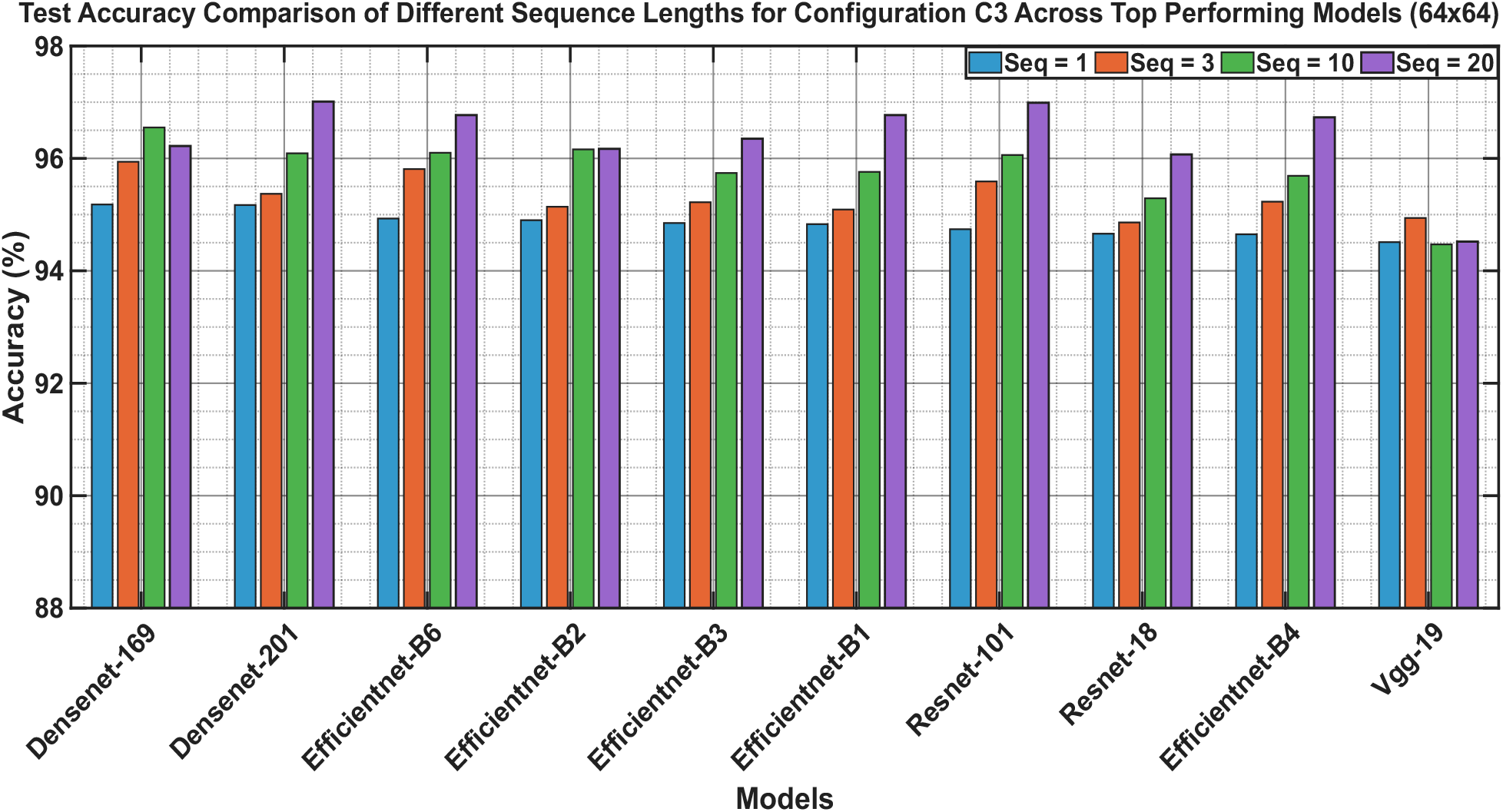
Comparison of Overall Accuracy Across input sequence length 1-20 for the top 10 Performing Models (64 *×* 64).

For DenseNet-201, the classification accuracy improved from 95.17% for a single frame to 97.01% for sequence length 20 and the ROC-AUC was improved from 0.9864 to 0.9943. Also, EfficientNet-B6 achieved 96.77% of accuracy with ROC-AUC of 0.9929 for sequence length 20. Similar performance improvements were observed for DenseNet, ResNet, HRNet, DeepLabV3, and Attention U-Net architectures. These results confirm that temporal dynamics carry relevant biological information related to cell spreading, activation, clustering and proliferative progression that cannot be fully extracted from static single-frame representations.

The results also show that the classification performance significantly improves until the sequence length of 10. But performance gains are marginal for most architectures after this point, indicating performance saturation. This suggests that the temporal features required to discriminate between the dormant and proliferative states are mostly present in the first 10 temporal observations. Increasing the length of the sequence beyond this point adds computational complexity but only limited improvement in predictive performance.

Across multiple temporal configurations, the highest classification performance was consistently demonstrated by DenseNet-201, EfficientNet-B6, DenseNet-169, EfficientNetB3, and EfficientNet-B7 among all evaluated models. These models obtained test accuracies *>*96% and ROC-AUC values *>*0.99 for longer temporal sequences, indicating their strong ability to capture complex spatio-temporal cellular dynamics. The confusion matrix and ROC for DenseNet-169, DenseNet201, EfficientNet-B7, EfficientNet-B2, and ViT for sequence lengths 1, 3, 10, and 20 are shown in Fig. 4. Overall, the experimental analysis demonstrates that both transferlearning strategy and temporal sequence modeling play crucial roles in automated dormant versus proliferative cancer-cell classification. The results highlight the importance of spatiotemporal representation learning and establish DenseNet and EfficientNet-based architectures as highly effective frameworks for analyzing dynamic cancer-cell behavior in physiologically relevant microenvironments. The corresponding sequence-length analysis for Configurations C1, C2, and C4 is provided in the supplementary material (Table S1, S2 and S3).

**Fig. 4:**
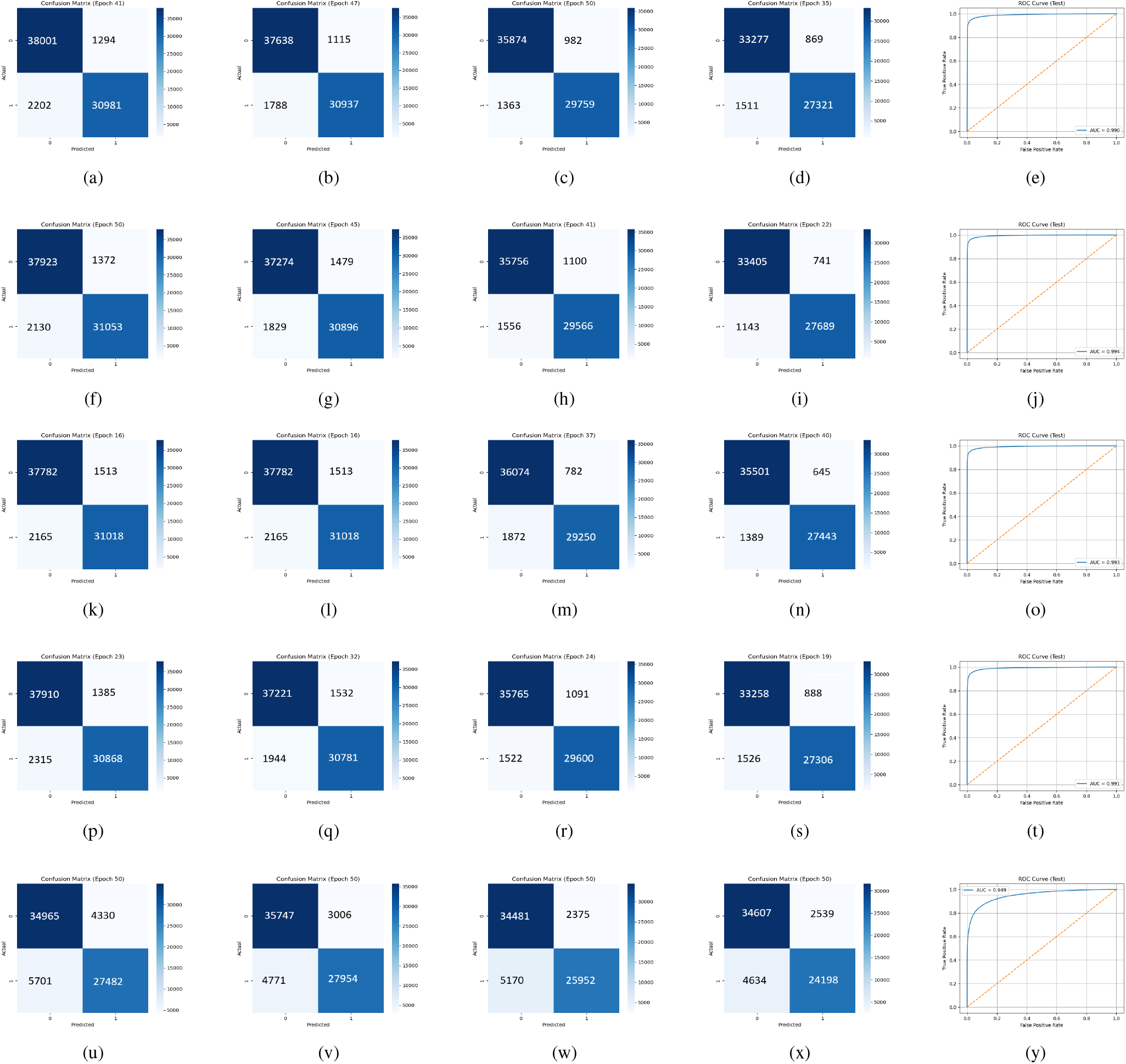
For DenseNet-169, Figures 4(a)–4(d) present the confusion matrices for sequence lengths 1, 3, 10, and 20, respectively, while Fig.4(e) shows the ROC curve for sequence length 20. Similarly, Figures 4(k)–4(o), Figures 4(p)–4(t), and Figures 4(u)–4(y) present the confusion matrices and ROC curves for DenseNet-201, EfficientNet-B7, EfficientNet-B2, and ViT, respectively, for the same sequence lengths.

In the next section we investigated the effect of input image size by increasing the resolution from 64*×* 64 to 96*×* 96. We hypothesize that the input size of 64×64 might be insufficient to capture all the spatial information given the size of the cells, their density and the impact of neighboring structures and contextual factors.

### C. *Effect of Temporal Sequence Length on* 96 *×* 96 *Images*

In this section, we set the input image size to 96*×* 96 and use training Configuration C3, which showed the best performance for the 64*×* 64 image setting previously. Here we analyze the model performance on 96*×* 96 inputs with different input sequence length from 1 to 20 to study the effect of temporal information on the model performance.

Table IV shows the effect of the temporal sequence length on the classification performance for the 96 *×* 96 image resolution in configuration C3. Among all the models investigated, EfficientNet-B7 performed especially well for longer temporal sequences with 98.86% accuracy and the highest ROC-AUC of 0.9982. EfficientNet-B6 showed one of the best overall performances with accuracies of 98.08%, 98.25%, 98.44%, and 98.66% for sequence lengths 1, 3, 10, and 20, respectively, with the ROC-AUC increasing from 0.9973 to 0.9989. Similarly, DenseNet-169 attained accuracy of 97.86% to 98.69% with ROC-AUC greater than 0.998 while DenseNet-201 attained highest accuracy of 98.70% and ROC-AUC of 0.9989.

The results also indicate that EfficientNet and DenseNet families consistently outperform competing architectures for all temporal configurations. The ResNet based architectures also showed highly competitive performance with ResNet-50 achieving 98.56% accuracy for sequence length 10 and ResNet-101 achieving more than 98.5% accuracy for longer temporal sequences. Among the segmentation based archi-tectures, DeepLabV3, Attention U-Net, HRNet, U-Net and U-Net++ achieved classification accuracy between 97% and 98.3%, indicating that segmentation based feature extraction could also provide highly discriminative representation for cancer-cell classification.

Fig. 5 compares the best performing models for different temporal sequence lengths, which emphasizes the importance of temporal information for classification accuracy. The overall trend of improved classification accuracy with increased temporal sequence length is consistent with the 64 *×*64 experiments. Most architectures showed significant performance gains when the sequence length was increased from 1 to 10 frames. However, the performance improvement above sequence length 10 became relatively smaller, suggesting a saturation effect. For instance, as the sequence length increases from 10 to 20, EfficientNet-B6 increases from 98.44% to 98.66% and DenseNet-169 increases from 98.35% to 98.69%. Similarly, DenseNet-201 improved from 97.82% to 98.65%, as sequence length increased from 1 to 10. However, a small improvement is observed in accuracy from 98.65% to 98.7% while increasing sequence length 10 to 20. Similar trends are observed for ResNet and DenseNet architecture. These results suggest that most of the discriminative temporal information is available in the first 10 observations with longer sequences providing marginal benefits.

**Fig. 5:**
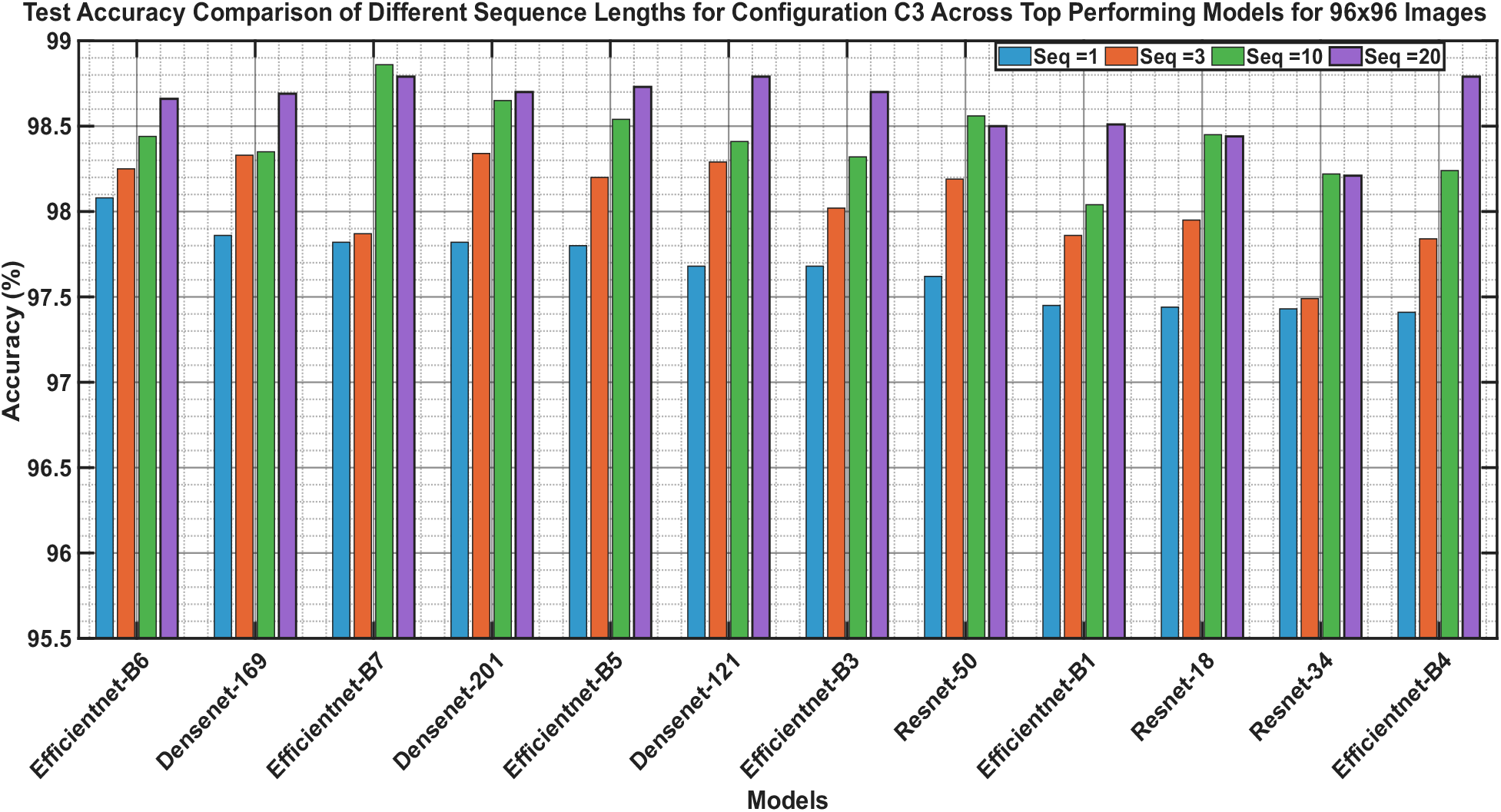
Comparison of Overall Accuracy Across input sequence length 1-20 for the top 10 Performing Models on 96 *×*96

Overall, the results indicate that the best representation for analyzing cancer-cell behavior is a combination of larger spatial coverage and temporal sequence modeling. The superior results of EfficientNet-B6, EfficientNet-B7, DenseNet-169, and DenseNet-201 also reflect the effectiveness of deep convolutional feature extraction to capture the complex spatiotemporal patterns in physiologically relevant breast cancercell microenvironments. The confusion matrix and ROC for EfficientNet-B6, DenseNet-169, EfficientNet-B7, DenseNet-201, and ViT for sequence lengths 1, 3, 10, and 20 are shown in Fig. 6. Furthermore, the corresponding results for Configurations C1, C2, and C4 are provided in the supplementary material (Table S4, S5, S6, and S7).

**Fig. 6:**
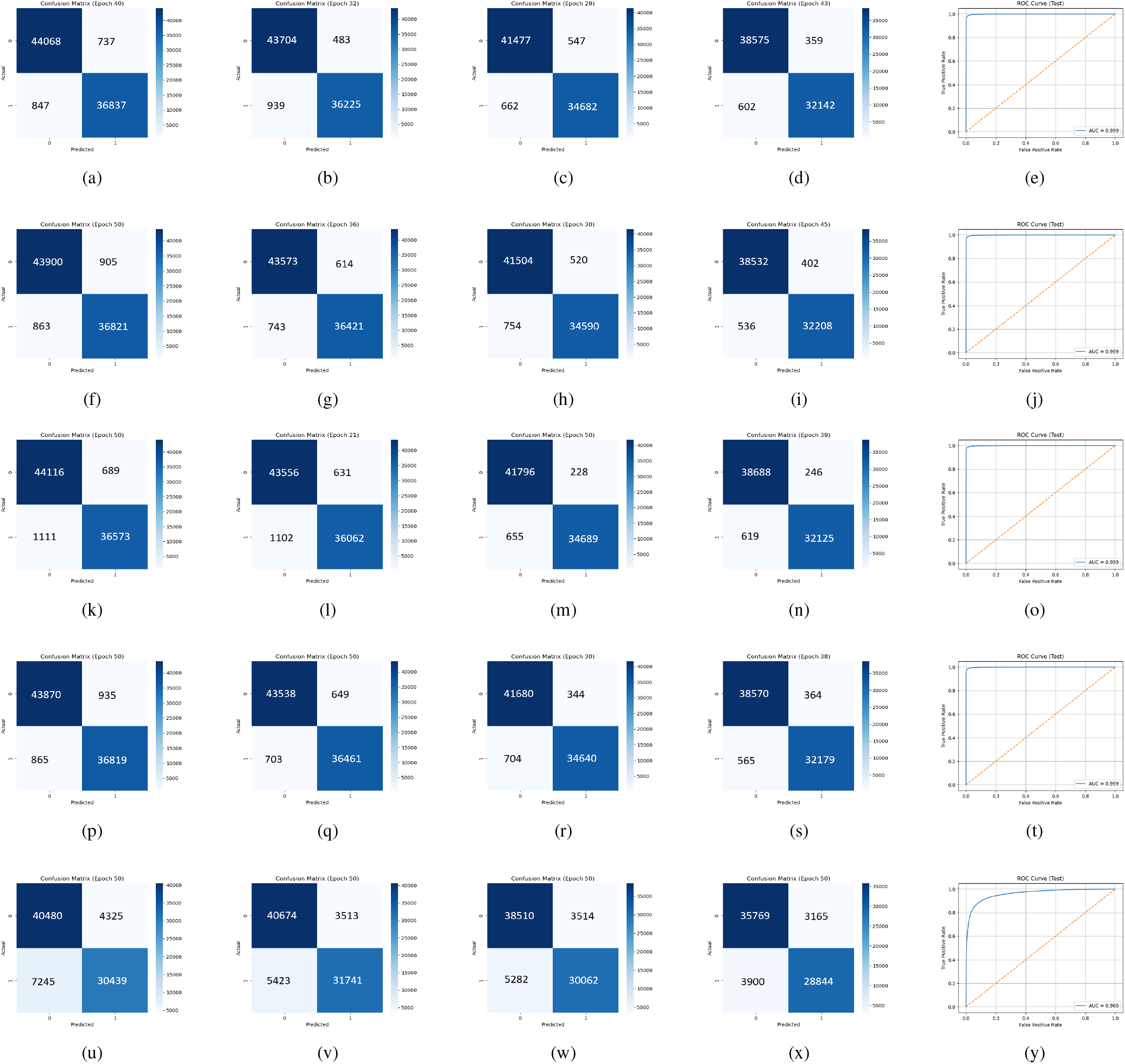
For EfficientNet-B6, Figures 6(a)–6(d) present the confusion matrices for sequence lengths 1, 3, 10, and 20, respectively, while Figure 6(e) shows the ROC curve for sequence length 20. Similarly, Figures 6(k)–6(o), Figures 6(p)–6(t), and Figures 6(u)–6(y) present the confusion matrices and ROC curves for DenseNet-169, EfficientNet-B7, DenseNet-201, and ViT, respectively, for the same sequence lengths.

The results obtained for the two image resolutions 64 *×*64 and 96 *×*96 indicate that classification performance is influenced by both temporal dynamics and spatial context. Therefore, the following subsection provides a combined analysis to evaluate the individual and joint effects of image resolution and temporal sequence length on dormant versus proliferative cancer-cell classification.

### D. Combined Analysis of Spatial Resolution and Temporal Information

The characterization of dormant and proliferative cancer cells depends on both spatial morphological information and temporal cellular dynamics. While the previous subsections examined these factors separately, the following analysis investigates their combined influence on classification performance and identifies the most effective spatio-temporal representation for cancer-cell behavior analysis. The Fig. 7 presents the classification performance of the best performing architectures for different temporal sequence lengths and image resolutions. The results indicate that both temporal information and spatial context are important for the accurate classification of dormant and proliferative breast cancer cells.

**Fig. 7:**
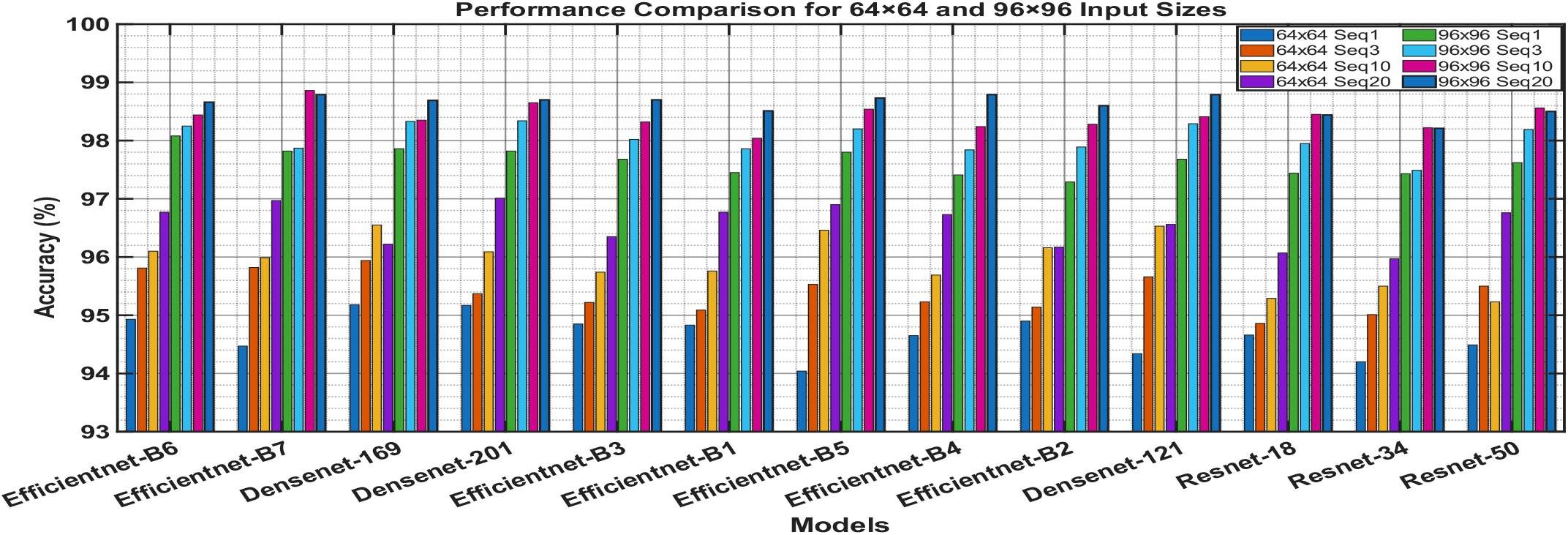
Effect of Sequence Length and Spatial Resolution

For all architectures evaluated, classification performance generally tended to increase with increasing sequence length. This trend was consistently observed for both 64*×* 64 and 96*×* 96 input resolutions, highlighting the significance of temporal modeling to capture dynamic cellular behaviours such as spreading, clustering and proliferative progression. The improvement was particularly visible when the sequence length was increased from 1 to 10 frames. However, for most models, only marginal gains were observed beyond sequence length 10, indicating a saturation effect. For example, when the sequence length was increased from 10 to 20, the accuracy of EfficientNet-B6 was increased from 98.44% to 98.66%, and the accuracy of DenseNet-169 was increased from 98.35% to 98.69%. We observed a similar behavior for DenseNet, EfficientNet and ResNet architectures, which means that the most discriminative temporal information is captured within the first 10 observations.

In addition to temporal information, the results demonstrate that increasing the spatial resolution from 64*×* 64 to 96 *×* 96 consistently improves classification performance across nearly all architectures and temporal configurations. The larger spatial region preserves additional cellular morphology, local structural characteristics, and surrounding microenvironmental information that may be lost in smaller image crops. Consequently, the networks are able to learn richer spatial representations associated with dormant and proliferative cellular states.

The performance gain achieved by increasing the image resolution is significant. EfficientNet-B6 achieved 96.77% accuracy with 64 × 64 images and sequence length 20, while the same architecture achieved 98.66% accuracy with 96*×* 96 images, which is an increase of about 1.9 percentage points. Similarly, DenseNet-201 was improved from 97.01% to 98.70% and DenseNet-169 was improved from 96.22% to 98.69% under the same temporal configuration. Most EfficientNet, DenseNet and ResNet architectures showed similar improvements, demonstrating that larger spatial coverage provides valuable contextual information for cancer-cell characterization.

For both image resolutions and different temporal configurations, the architectures DenseNet-169, DenseNet-201 and EfficientNet-B6 obtained the best performance. DenseNet-169 and DenseNet-201 achived max classification accuracy 98.69% and 98.70% respectively. EfficientNet-B6 demonstrated exceptional performance, with a classification accuracy of 98.66% and a ROC-AUC of 0.9989, and EfficientNet-B7 was the best with a ROC-AUC of 0.9990.

A comparison of the experimental results further reveals that spatial resolution has a greater influence on classification performance than temporal sequence length. Increasing the image resolution from 64 *×* 64 to 96*×* 96 resulted in an improvement of approximately 2–4 percentage points for the top-performing models, whereas increasing the sequence length from a single frame to 20 frames typically improved performance by approximately 1–2 percentage points. These findings suggest that spatial morphological characteristics and the surrounding cellular microenvironment provide more discriminative information than long temporal sequences alone. Temporal information is still useful though, as it consistently improves performance for almost all architectures and resolutions.

Lastly, the experimental analysis shows that the best representation is obtained by exploiting both larger spatial contexts and temporal sequence information. The spatial resolution provides a significant performance benefit however the temporal modelling provides additional information that further improves the classification accuracy. These results emphasize the need to model the morphology and temporal evolution of cells simultaneously and indicate that EfficientNet and DenseNet architectures are appropriate frameworks for the automated analysis of dormant and proliferative breast cancercell behavior in physiologically relevant microenvironments.

### E. Why Do Some Models Perform Better? An Analysis of Early-Stage Cellular-State Discrimination

Besides overall classification accuracy and ROC-AUC, it is also important to evaluate how the DL model performance evolves over the temporal sequence, as considered in the previous analyses. Early frames in particular are often more difficult since dormant and proliferative cells have similar morphological characteristics during the first stages of observation. Hence, the capacity to correctly classify cells at early time points provides additional insight into the discriminative power of different architectures.

The Fig. 8 shows the accuracy comparison between the worst and the best models per frame and per class for different sequence length and input resolution. The figure compares the performance of the ViT and EfficientNet-B6 architectures for sequence lengths 1 and 20 frames and image resolutions 64×64 and 96×96 pixels. The results show large differences in their ability to distinguish between dormant and outbreaking cell states in the early stages of the sequence, with EfficientNet-B6 generally more stable and higher classification accuracy across frames than the corresponding ViT models.

**Fig. 8:**
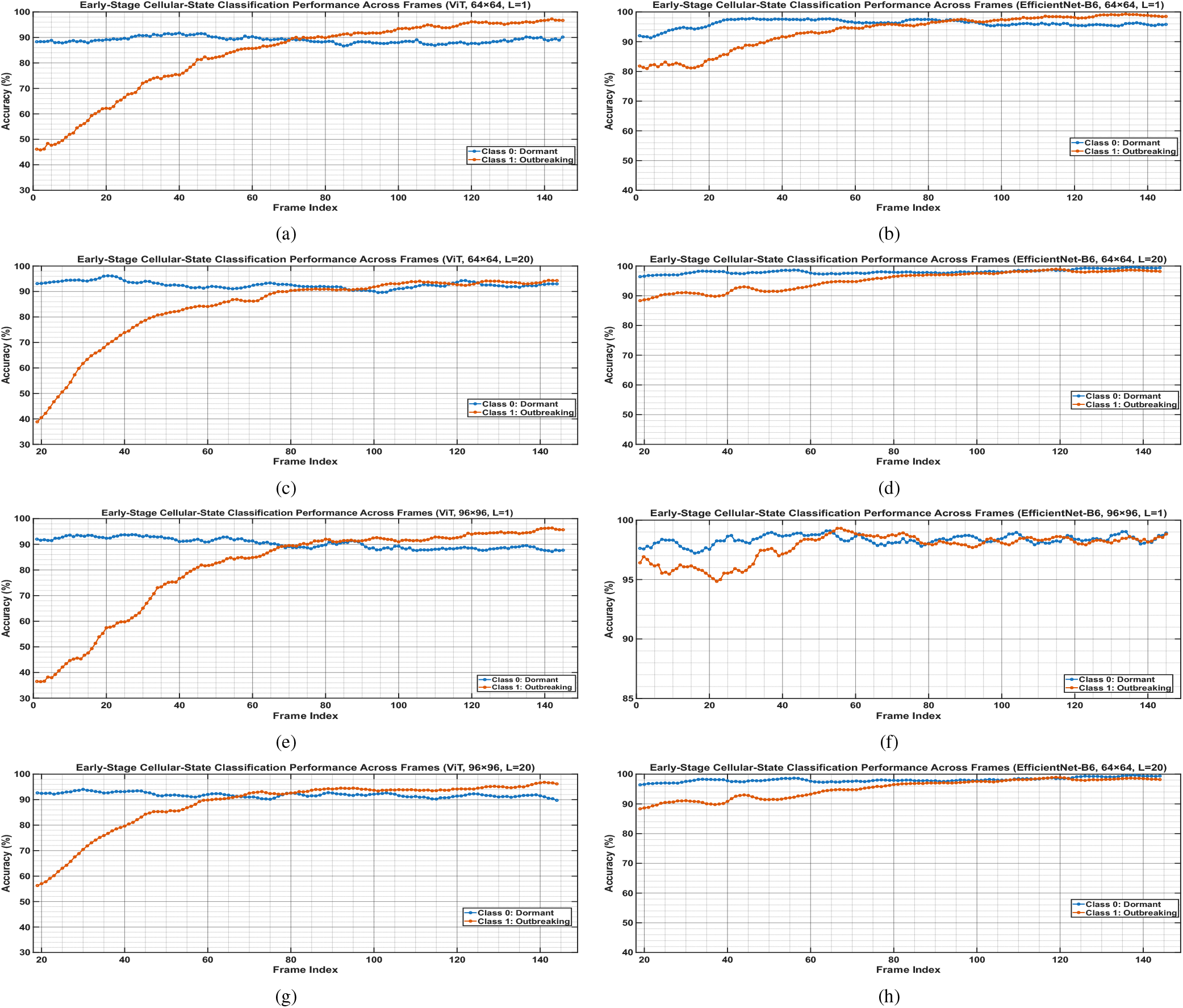
Frame-wise and class-wise accuracy comparison between the worst-performing and best-performing models. Figures 8(a) and 8(b) present the results for sequence length 1 with input sizes of 64×64 using ViT and EfficientNet-B6, respectively. Figures 8(c) and 8(d) show the corresponding results for sequence length 20 with an input size of 64×64. Figures 8(e) and 8(f) illustrate the performance for sequence length 1 with an input size of 96×96 using ViT and EfficientNet-B6, respectively. Finally, Figures 8(g) and 8(h) depict the results for sequence length 20 with an input size of 96×96 using ViT and EfficientNet-B6, respectively.

The frame-wise class-wise accuracy curves of ViT exhibit a large variance in all experimental settings. Class 1 (Outbreaking) accuracy is significantly lower in the early frames, often starting between 40-60% and gradually increasing as more temporal information is acquired, in both the 64×64 and 96×96 input resolutions. On the other hand, Class 0 (Dormant) shows relatively high accuracy along the sequence, typically higher than 85%. This difference indicates that the ViT model has difficulty in correctly identifying the outbreaking cellular state during the early stage of observation, where morphological difference between dormant and proliferative cells is still subtle, as also shown in Fig. 1. The Class 1 accuracy rises smoothly over later frames, finally reaching a performance close to Class 0, but the early-stage classification gap persists for both sequence lengths and image resolutions.

In contrast, EfficientNet-B6 achieves consistently high performance across all experiment setups. The accuracies for the dormant and outbreaking classes stay high over the entire frame sequence for both sequence lengths (1 and 20) as well as for both input resolutions (64×64 and 96×96). The class-wise accuracy curves are very close to each other and mostly above 90-95% with only minor fluctuations during the earliest frames. Interestingly, the 96×96 configurations show near-perfect and highly stable performance on both classes, indicating that EfficientNet-B6 can extract highly discriminative spatial features even when phenotypic differences between cellular states are limited.

For the ViT model, increasing the sequence length from 1 to 20 yielded only marginal improvements, with a slight decrease in the class-wise accuracy gap at later frames. However the challenge of correctly identifying outbreaking cells in the early stages persists. On the contrary, EfficientNet-B6 consistently achieved high and balanced accuracies for both sequence lengths, indicating that the performance is primarily driven by effective spatial feature extraction rather than dependence on extended temporal information.

These results also justify the overall better performance of EfficientNet-B6 and other best performing models like dence-169 and DenseNet-201. It is more accurate on average, and it can reliably classify the two cellular states along the entire sequence, including the most difficult early frames, leveraging the higher average accuracy. Biologically, the results show that convolutional architectures are more appropriate to learn subtle spatial features related to early transitions between cellular states, enabling reliable discrimination before major phenotypic divergence. This capability is particularly important for automated analysis of cancer cell dormancy and metastatic progression where a precise identification of proliferative activation at the earliest observable stages may provide substantial insights to disease development and therapeutic intervention.

### F. Statistical Significance Analysis

To further evaluate the robustness and reproducibility of the experimental results, a further statistical significance analysis was performed. To further assess the robustness and repro-ducibility of the experimental findings, an additional statistical significance analysis was conducted. Rather than evaluating all experimental settings, a representative configuration was selected to provide a focused assessment of performance variability and model stability. Specifically, the analysis was performed using the 96 *×*96 image resolution, transferlearning configuration C3, and sequence length 1. The selected configuration combines the best-performing image resolution and transfer-learning strategy identified in the previous experiments, while the shortest sequence length was adopted to reduce computational complexity during repeated training runs. This setting therefore provides an effective benchmark for evaluating the statistical significance of performance differences among the competing architectures.

Table V shows the performance of all the investigated architectures under the selected experimental configuration on 10 independent runs. The results show that EfficientNet-B7 achieved the highest mean accuracy of 97.842% with a standard deviation of 0.3287, followed by EfficientNet-B5 (97.807%), EfficientNet-B6 (97.800%), DenseNet-201 (97.782%) and DenseNet-169 (97.755%). The difference in the average accuracy between the top models is not large, but it is important to consider the stability of the performance over repeated runs to evaluate the robustness of the models.

**TABLE IV:**
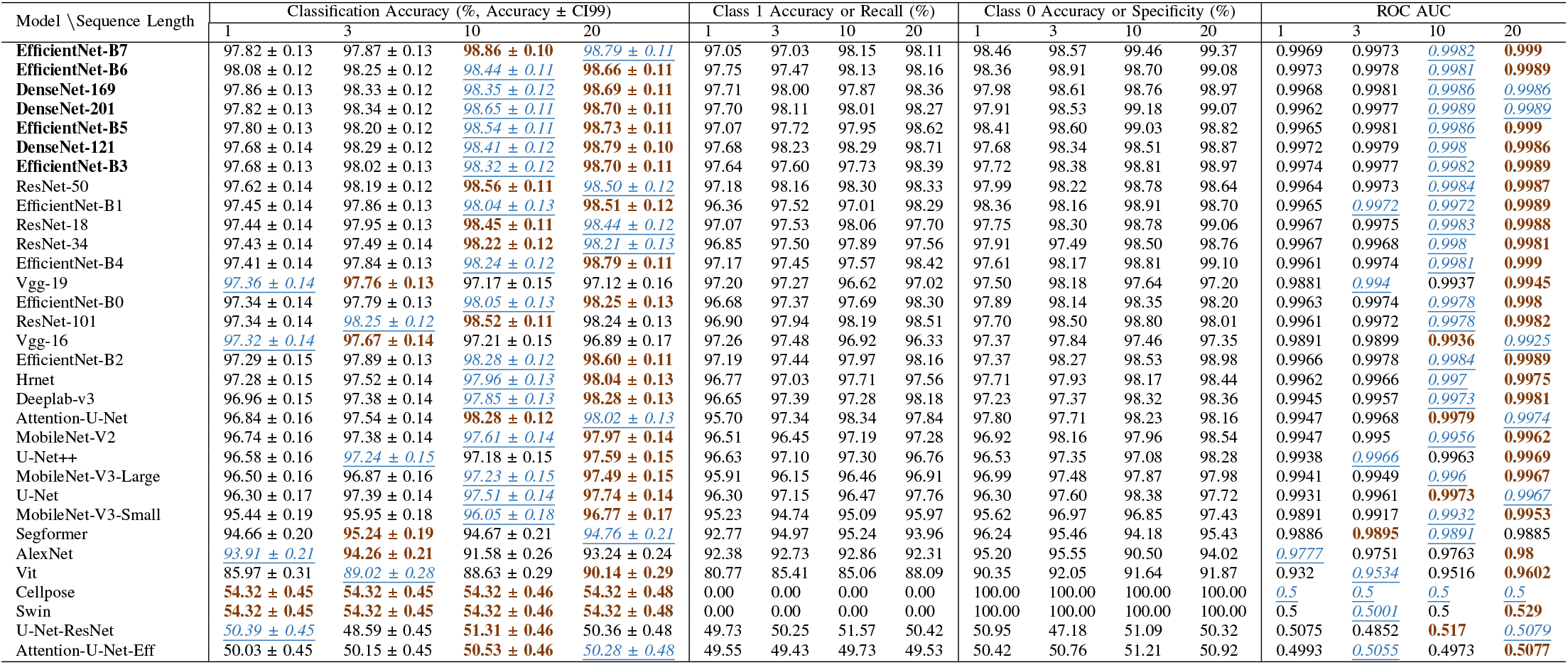
Performance Comparison of Advanced DL Models Based on Classification Accuracy and ROC Metrics Across Sequence Lengths 1, 3, 10, and 20 for 96 *×*96 Input Images. The best and second-best configurations for each model are highlighted in **bold** and *<u>italicized-underlined</u>* text, respectively.

**TABLE V:** Performance comparison of all evaluated models across 10 independent runs. Mean accuracy, standard deviation, and per-run test accuracies are reported.

| Model | MeanAccuracy | StdAccuracy | Test Acc (%) |  |  |  |  |  |  |  |  |  |
| --- | --- | --- | --- | --- | --- | --- | --- | --- | --- | --- | --- | --- |
|  |  |  | Run 1 | Run 2 | Run 3 | Run 4 | Run 5 | Run 6 | Run 7 | Run 8 | Run 9 | Run 10 |
| **EfficientNet-B7** | **97.842** | 0.3287 | 98.148 | **98.460** | 98.103 | 97.431 | 97.527 | 97.751 | 97.831 | 97.453 | 97.872 | 97.844 |
| **EfficientNet-B5** | 97.807 | 0.2810 | 97.920 | 98.323 | 98.032 | 97.425 | 97.456 | 97.604 | 97.664 | 97.737 | 97.952 | 97.962 |
| **EfficientNet-B6** | 97.800 | 0.3289 | 98.219 | 98.219 | 98.007 | 97.285 | 97.289 | 97.635 | 97.815 | 97.727 | 97.927 | 97.873 |
| **DenseNet-201** | 97.782 | 0.3324 | 97.935 | 98.046 | 98.243 | 97.123 | 97.620 | 97.444 | 97.630 | 97.960 | 98.026 | 97.794 |
| **DenseNet-169** | 97.755 | 0.2781 | 97.812 | 97.989 | 98.205 | 97.330 | 97.565 | 97.467 | 98.062 | 97.847 | 97.588 | 97.686 |
| DenseNet-121 | 97.615 | 0.2545 | 97.609 | 97.845 | 98.030 | 97.386 | 97.372 | 97.161 | 97.771 | 97.729 | 97.589 | 97.662 |
| EfficientNet-B3 | 97.438 | 0.2549 | 97.597 | 97.765 | 97.640 | 97.265 | 97.047 | 97.419 | 97.064 | 97.591 | 97.666 | 97.324 |
| EfficientNet-B2 | 97.429 | 0.3695 | 97.612 | 98.013 | 97.747 | 96.979 | 96.935 | 97.541 | 97.072 | 97.391 | 97.777 | 97.218 |
| ResNet-50 | 97.422 | 0.3140 | 97.580 | 97.438 | 97.985 | 97.202 | 97.359 | 97.281 | 97.624 | 97.467 | 96.773 | 97.517 |
| EfficientNet-B1 | 97.388 | 0.3052 | 97.560 | 98.046 | 97.560 | 97.164 | 97.074 | 97.208 | 97.056 | 97.251 | 97.576 | 97.388 |
| ResNet-101 | 97.330 | 0.3296 | 97.171 | 97.778 | 97.681 | 96.735 | 97.367 | 97.081 | 97.718 | 97.341 | 97.086 | 97.340 |
| EfficientNet-B4 | 97.305 | 0.3792 | 97.648 | 97.929 | 97.775 | 96.839 | 96.984 | 97.061 | 97.212 | 96.910 | 97.444 | 97.249 |
| ResNet-18 | 97.287 | 0.3021 | 97.419 | 97.899 | 97.418 | 97.001 | 97.051 | 97.009 | 97.270 | 97.556 | 97.313 | 96.935 |
| ResNet-34 | 97.284 | 0.3273 | 97.322 | 97.903 | 97.560 | 96.897 | 97.097 | 97.055 | 97.326 | 97.130 | 97.623 | 96.930 |
| Vgg-19 | 97.148 | 0.3585 | 97.183 | 97.712 | 97.617 | 96.701 | 96.849 | 96.784 | 97.422 | 97.305 | 96.852 | 97.055 |
| Vgg-16 | 97.111 | 0.3579 | 97.303 | 97.617 | 97.457 | 96.331 | 96.957 | 96.987 | 97.319 | 96.997 | 97.196 | 96.939 |
| EfficientNet-B0 | 97.095 | 0.3511 | 97.255 | 97.863 | 97.296 | 96.616 | 96.691 | 97.010 | 96.996 | 96.909 | 97.184 | 97.125 |
| Hrnet | 96.951 | 0.4249 | 97.118 | 97.056 | 97.589 | 96.298 | 96.888 | 96.173 | 97.068 | 97.280 | 97.097 | 96.941 |
| Attention-U-Net | 96.776 | 0.4012 | 97.006 | 97.485 | 97.233 | 96.166 | 96.685 | 96.428 | 96.691 | 96.769 | 96.914 | 96.379 |
| MobileNet-V2 | 96.691 | 0.4013 | 96.497 | 97.342 | 97.234 | 96.303 | 96.512 | 96.072 | 96.492 | 96.820 | 96.957 | 96.684 |
| DeepLab-v3 | 96.664 | 0.3193 | 97.037 | 97.095 | 96.938 | 96.044 | 96.695 | 96.359 | 96.738 | 96.545 | 96.664 | 96.524 |
| U-Net++ | 96.616 | 0.5358 | 96.671 | 97.615 | 97.183 | 95.977 | 96.174 | 96.317 | 96.116 | 96.539 | 97.141 | 96.423 |
| U-Net | 96.589 | 0.5182 | 96.550 | 97.245 | 97.197 | 95.655 | 96.579 | 96.308 | 96.159 | 96.791 | 97.157 | 96.249 |
| MobileNet-V3-Large | 96.075 | 0.3826 | 96.246 | 96.807 | 96.226 | 95.563 | 95.654 | 95.934 | 96.383 | 95.780 | 95.871 | 96.288 |
| MobileNet-V3-Small | 95.426 | 0.4329 | 96.238 | 95.733 | 95.758 | 95.052 | 95.289 | 94.987 | 94.885 | 95.504 | 95.699 | 95.113 |
| Segformer | 94.707 | 0.5429 | 94.722 | 95.569 | 95.295 | 93.627 | 94.641 | 94.338 | 94.375 | 94.835 | 95.056 | 94.610 |
| AlexNet | 93.823 | 0.7147 | 93.896 | 94.817 | 95.228 | 92.875 | 93.647 | 93.965 | 93.181 | 93.446 | 93.513 | 93.658 |
| Vit | 86.754 | 1.3683 | 84.153 | 86.556 | 88.013 | 87.202 | 88.586 | 86.780 | 86.173 | 87.776 | 84.968 | 87.330 |
| Cellpose | 54.316 | 0.0000 | 54.316 | 54.316 | 54.316 | 54.316 | 54.316 | 54.316 | 54.316 | 54.316 | 54.316 | 54.316 |
| Swin | 54.316 | 0.0000 | 54.316 | 54.316 | 54.316 | 54.316 | 54.316 | 54.316 | 54.316 | 54.316 | 54.316 | 54.316 |
| Attention-U-Net-Eff | 50.275 | 0.5786 | 49.491 | 50.192 | 50.532 | 49.906 | 49.935 | 50.855 | 50.563 | 51.449 | 49.985 | 49.837 |
| U-Net-ResNet | 50.017 | 0.5747 | 50.470 | 49.496 | 50.343 | 50.441 | 50.203 | 49.514 | 49.095 | 50.205 | 49.503 | 50.897 |
| Mean on All models | 91.026 | 0.395 | 91.063 | 91.453 | 91.430 | 90.608 | 90.886 | 90.818 | 90.919 | 91.084 | 91.019 | 90.981 |
| Max. on All Models | **97.842** | 1.368 | 98.219 | **98.460** | 98.243 | 97.431 | 97.620 | 97.751 | 98.062 | 97.960 | 98.026 | 97.962 |

**TABLE VI:** Pairwise Wilcoxon signed-rank test *p*-values among the top-performing classification models. Values greater than 0.05 indicate that the performance difference between the corresponding model pair is not statistically significant, whereas values below 0.05 indicate a statistically significant difference.

|  | EfficientNet-B6 | DenseNet-169 | EfficientNet-B7 | DenseNet-201 | EfficientNet-B5 | DenseNet-121 | EfficientNet-B3 |
| --- | --- | --- | --- | --- | --- | --- | --- |
| **EfficientNet-B6** | NaN | $<10^{-4}$ | $<10^{-4}$ | $<10^{-4}$ | $<10^{-4}$ | $<10^{-4}$ | $<10^{-4}$ |
| **DenseNet-169** | $<10^{-4}$ | NaN | 0.006 | 0.095 | $<10^{-4}$ | $<10^{-4}$ | $<10^{-4}$ |
| **EfficientNet-B7** | $<10^{-4}$ | 0.006 | NaN | 0.108 | $<10^{-4}$ | $<10^{-4}$ | $<10^{-4}$ |
| **DenseNet-201** | $<10^{-4}$ | 0.095 | 0.108 | NaN | $<10^{-4}$ | $<10^{-4}$ | $<10^{-4}$ |
| **EfficientNet-B5** | $<10^{-4}$ | $<10^{-4}$ | $<10^{-4}$ | $<10^{-4}$ | NaN | $<10^{-4}$ | $<10^{-4}$ |
| **DenseNet-121** | $<10^{-4}$ | $<10^{-4}$ | $<10^{-4}$ | $<10^{-4}$ | $<10^{-4}$ | NaN | $<10^{-4}$ |
| **EfficientNet-B3** | $<10^{-4}$ | $<10^{-4}$ | $<10^{-4}$ | $<10^{-4}$ | $<10^{-4}$ | $<10^{-4}$ | NaN |

The results show that ranking of the models is highly consistent across all the 10 independent runs. Run 2 yielded slightly higher accuracies for most architectures, but the increase was small and did not influence the relative performance of models.

The small standard deviations and the stable performance across repeated experiments indicate good generalization capability and demonstrate that the training procedure is robust to random initialization and sampling effects. This stability is likely contributed by the adopted regularization strategies such as random data sampling and dropout during training.

To determine whether the observed differences between competing architectures are statistically meaningful, a pairwise Wilcoxon signed-rank test was performed using the accuracies obtained from the 10 independent runs. The complete pairwise significance matrix for all evaluated models is provided in Supplementary Table S8, while the statistical comparisons among the top-performing models are presented in Table VI. Results show that there is no statistically significant difference in performance between EfficientNet-B7, DenseNet-201, and DenseNet-169. Specifically, the comparison of EfficientNet-B7 versus DenseNet-201 resulted in a p-value of 0.108 and DenseNet-169 versus DenseNet-201 resulted in a p-value of 0.095. Both values are above the significance threshold of 0.05, therefore the null hypothesis cannot be rejected. This means similar performance of these models. For all other pairwise comparisons between the best performing architectures, the p-values were smaller than 10^*−*4^, suggesting statistically significant differences.

Therefore, although EfficientNet-B7 has the highest mean accuracy, the statistical analysis shows that its performance is not significantly different from DenseNet-201 and DenseNet-169. Therefore, we can consider these three architectures to be statistically equivalent under the evaluated experimental setting. EfficientNet-B7 provides the highest average performance among these three architectures.

## VI. Conclusion

This study presents a comprehensive evaluation of deep learning architectures for classification of dormant and proliferative breast cancer cells from microscopy image sequences. The experimental results showed that both spatial and temporal information is important for the classification performance and that larger image resolutions lead to larger improvements than longer temporal sequences. Temporal modeling enhanced the capability to capture dynamic cellular behavior; however, most of the performance improvements were observed with the 10 time frames.

Among the evaluated architectures, EfficientNet and DenseNet models consistently outperformed others with EfficientNet-B6, EfficientNet-B7, DenseNet-169 and DenseNet-201 achieving highest scores. Moreover, the transfer-learning analysis revealed that the partial finetuning of the higher-level network layers leads to a good compromise between the feature preservation and the domain adaptation. Incorporation of spatial and temporal information demonstrated that the best representation was obtained with the combined modeling of cellular morphology and temporal progression. The proposed framework achieved classification accuracies of almost 99% and ROC-AUCs of almost 1.0, showing the effectiveness of deep spatio-temporal learning for automated cancer-cell analysis using microscopy.

In summary, the results emphasize the importance of spatiotemporal representation learning for automated cancer-cell behavior analysis and demonstrate that deep learning can discriminate dormant vs. proliferative cellular states with high accuracy in physiologically relevant microenvironments. These results are promising for future AI-driven strategies for cancercell phenotyping, disease progression analysis, and evaluation of therapeutic response.

Future work will focus on hybrid architectures that combine the complementary advantages of the best performing EfficientNet and DenseNet models to further improve classification performance and robustness. Another promising direction involves the development of dedicated spatio-temporal frameworks that can explicitly model long-term cellular dynamics, such as recurrent neural networks, transformer-based architectures and state-space models. Additionally, the integration of cell-tracking data, multi-modal biological measurements, physics-informed neural networks and explainable artificial intelligence (XAI) techniques may provide more biological insights into the mechanisms that regulate cancer-cell dormancy, reactivation and metastatic progression. The generalizability of the proposed framework will be enhanced by applying it to larger datasets, multiple types of cancer and longitudinal studies and will contribute to the development of AI-assisted systems for cellular phenotyping, metastasis-risk assessment and prediction of therapeutic response.

## ACKNOWLEDGMENT

This work was supported by the Artificial Intelligence Research Center at the University of Haifa.

## Code Availability

The implementation of the proposed deep learning frameworks, along with all necessary scripts for training and evaluation, will be made publicly available. The source code can be accessed at: https://github.com/omveersharmanet/Deep Learning for Dormant and Outbreaking Metastatic Breast Tumor Cells A Benchmark Study.git

---

**TABLE S1:**
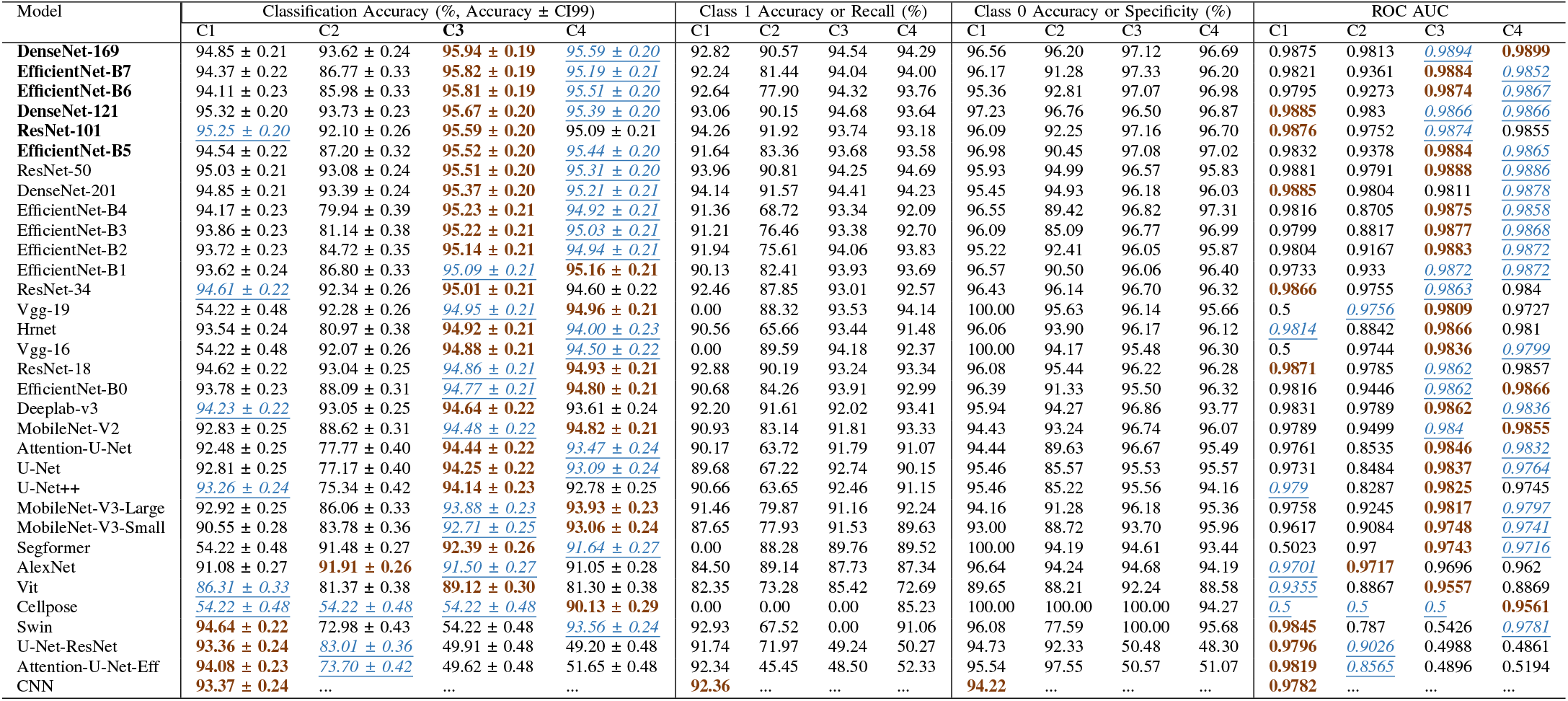
Performance comparison of advanced DL models for a sequence length of 3 and an input image size of 64*×* 64, evaluated using classification accuracy and ROC-AUC metrics under configurations C1, C2, C3, and C4. The best and second-best configurations for each model are highlighted in **bold** and *<u>italicized-underlined</u>* text, respectively.

**TABLE S2:**
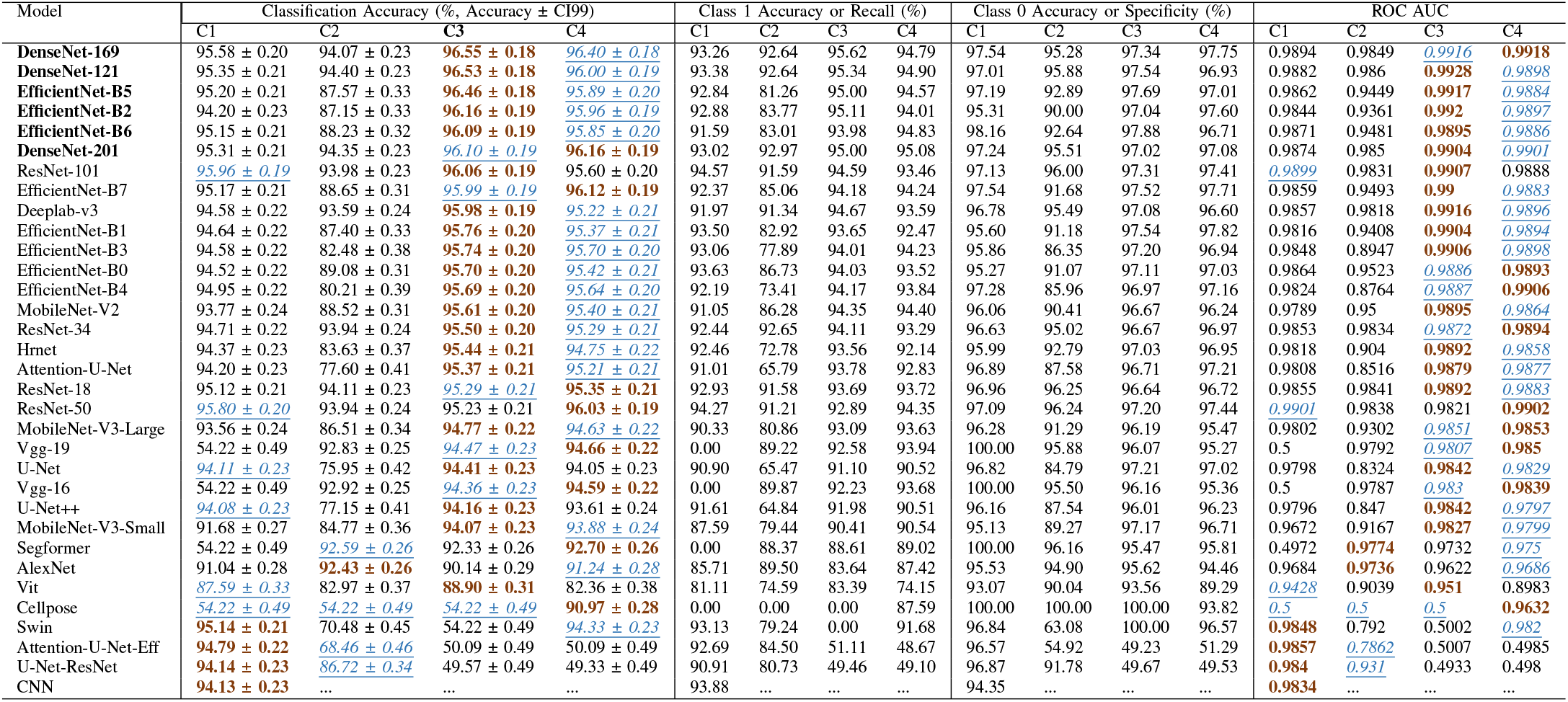
Performance comparison of advanced DL models for a sequence length of 10 and an input image size of 64 *×* 64, evaluated using classification accuracy and ROC-AUC metrics under configurations C1, C2, C3, and C4. The best and second-best configurations for each model are highlighted in **bold** and *<u>italicized-underlined</u>* text, respectively.

**TABLE S3:**
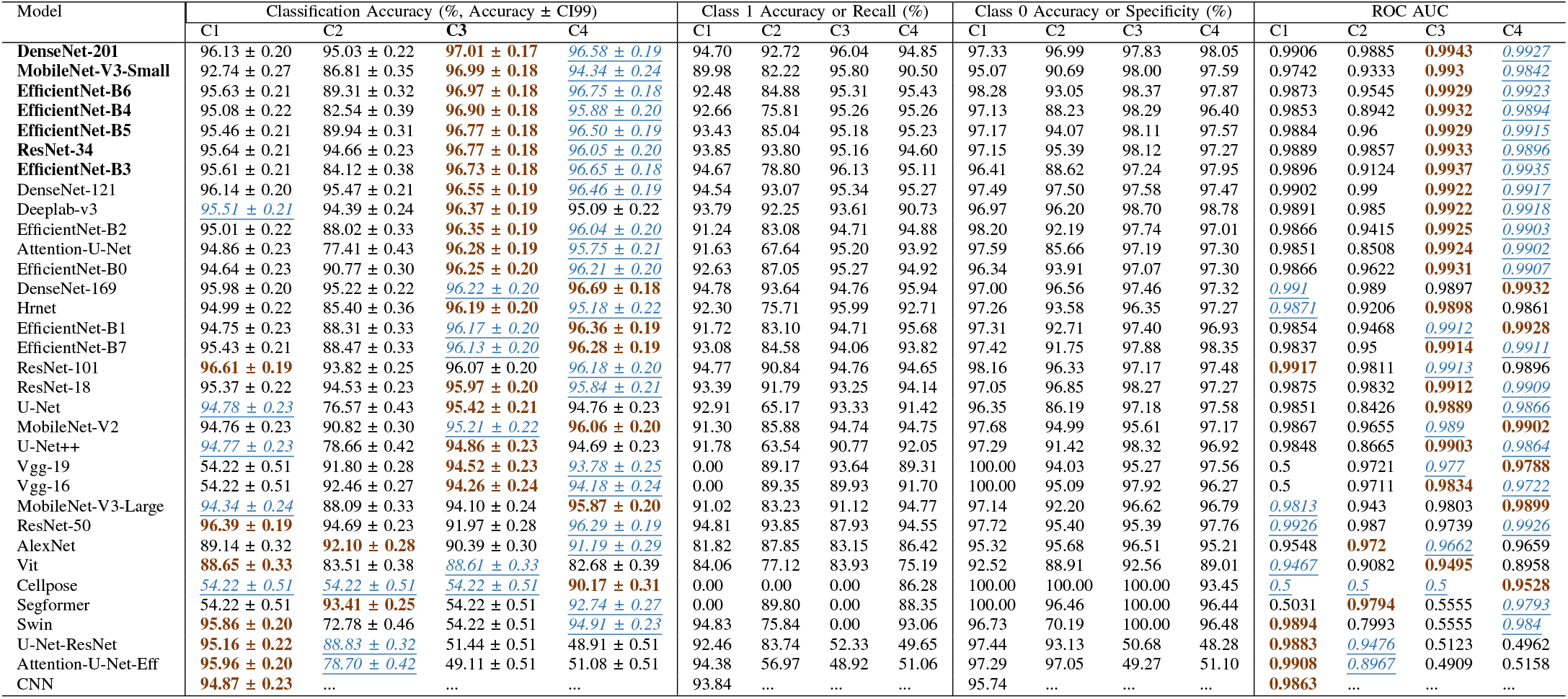
Performance comparison of advanced DL models for a sequence length of 20 and an input image size of 64 *×* 64, evaluated using classification accuracy and ROC-AUC metrics under configurations C1, C2, C3, and C4. The best and second-best configurations for each model are highlighted in **bold** and *<u>italicized-underlined</u>* text, respectively.

**TABLE S4:**
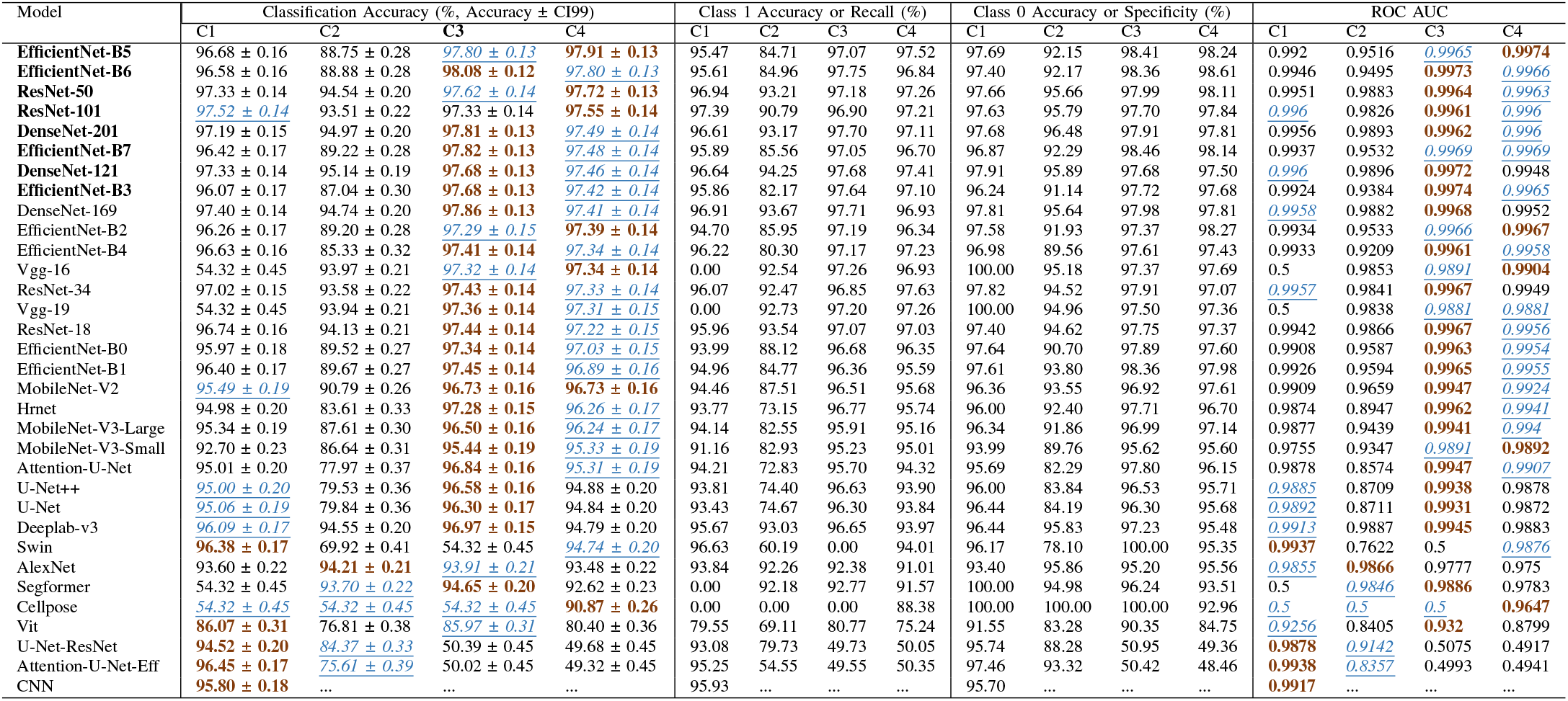
Performance comparison of advanced DL models for a sequence length of 1 and an input image size of 96 *×*96, evaluated using classification accuracy and ROC-AUC metrics under configurations C1, C2, C3, and C4. The best and second-best configurations for each model are highlighted in **bold** and *<u>italicized-underlined</u>* text, respectively.

**TABLE S5:**
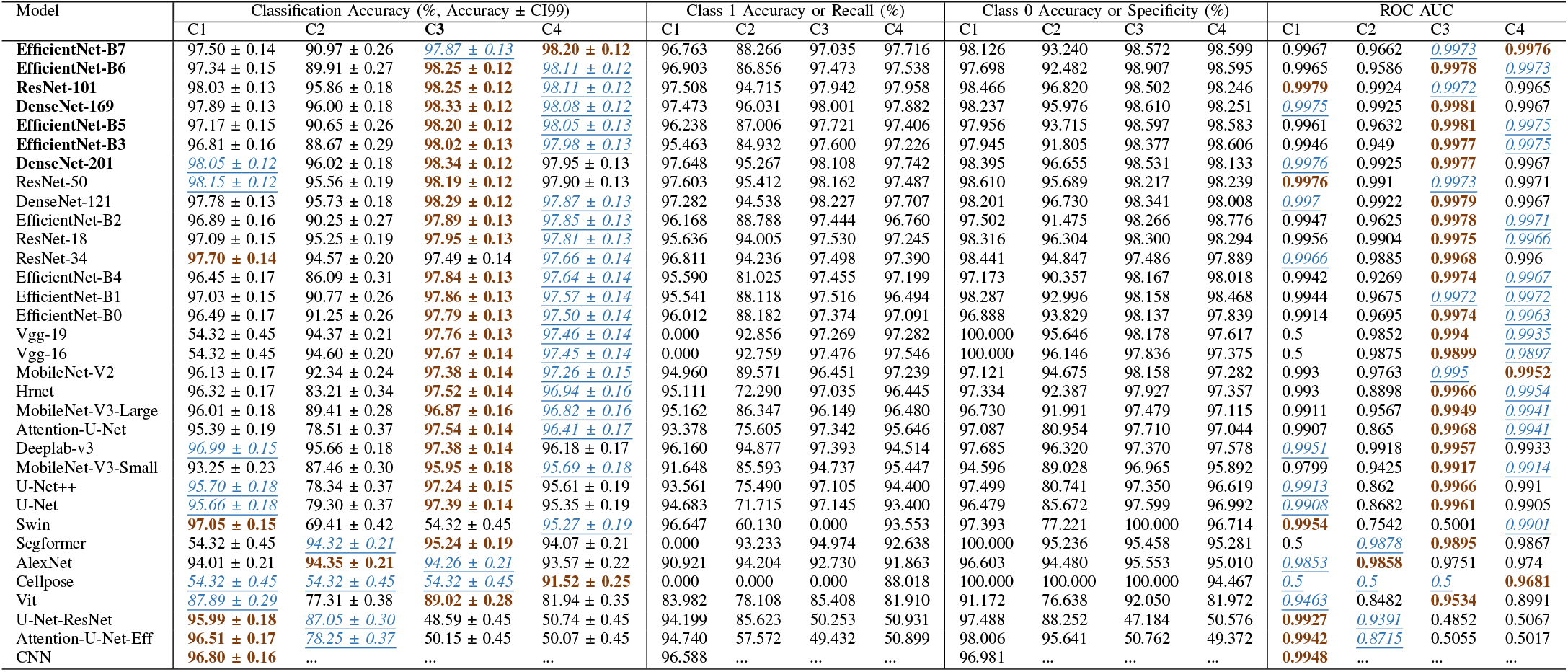
Performance comparison of advanced DL models for a sequence length of 3 and an input image size of 96 *×*96, evaluated using classification accuracy and ROC-AUC metrics under configurations C1, C2, C3, and C4. The best and second-best configurations for each model are highlighted in **bold** and *<u>italicized-underlined</u>* text, respectively.

**TABLE S6:**
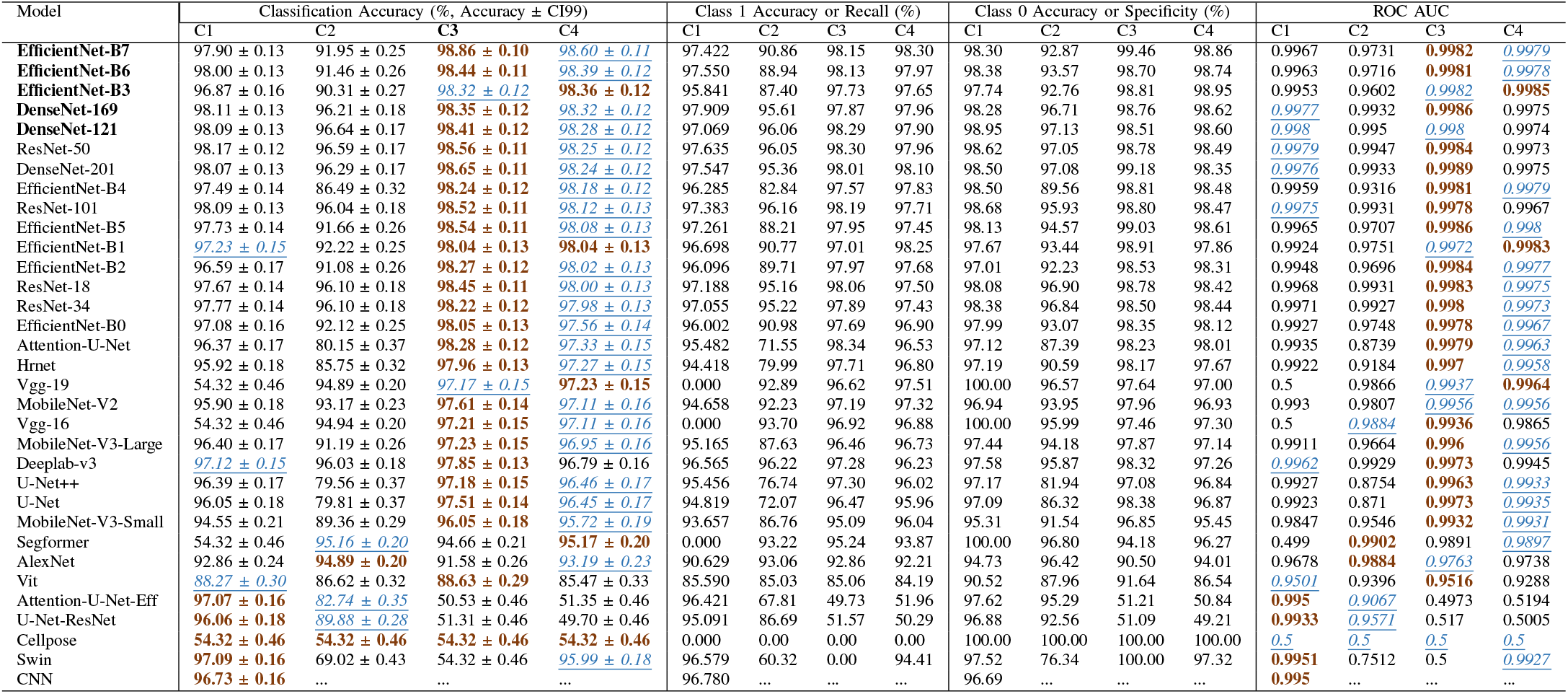
Performance comparison of advanced DL models for a sequence length of 10 and an input image size of 96 *×* 96, evaluated using classification accuracy and ROC-AUC metrics under configurations C1, C2, C3, and C4. The best and second-best configurations for each model are highlighted in **bold** and *<u>italicized-underlined</u>* text, respectively.

**TABLE S7:**
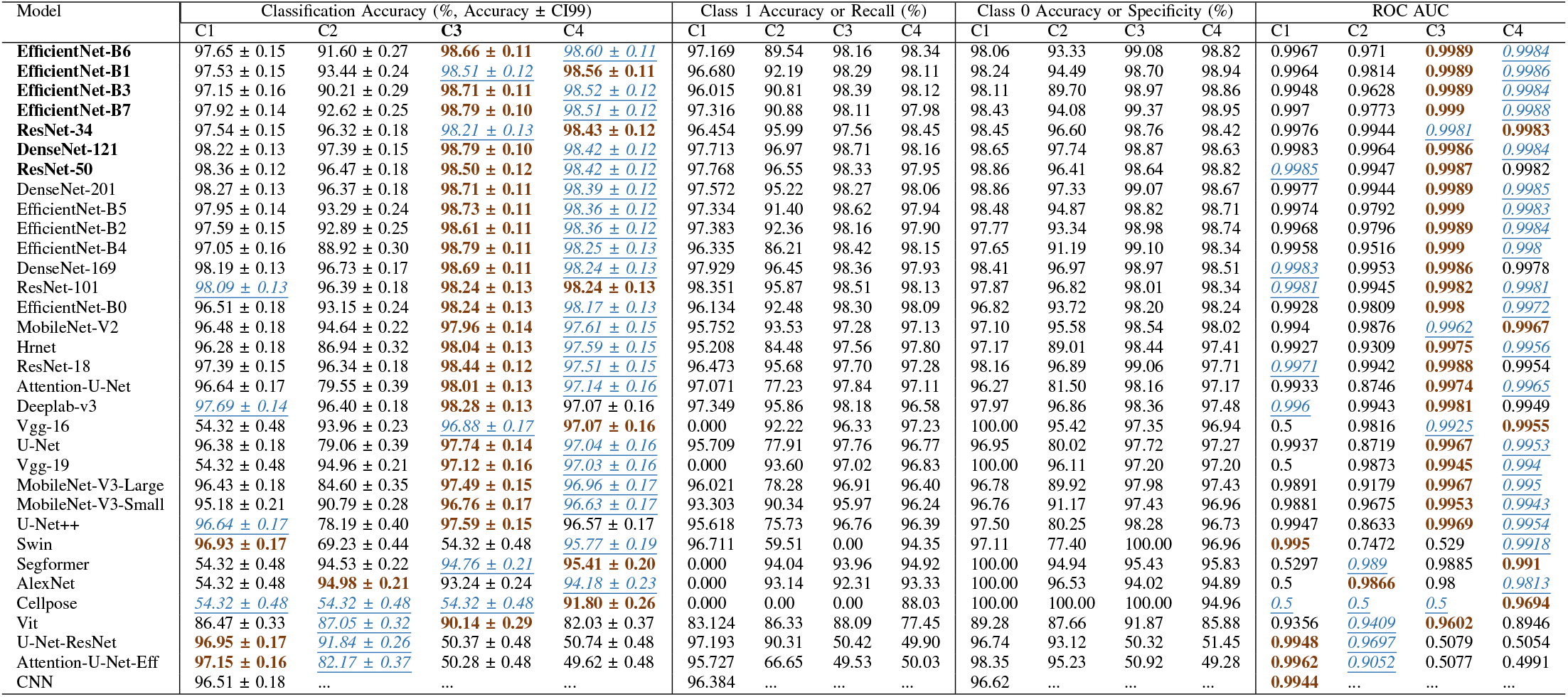
Performance comparison of advanced DL models for a sequence length of 20 and an input image size of 96 *×*96, evaluated using classification accuracy and ROC-AUC metrics under configurations C1, C2, C3, and C4. For each model, the best-performing configuration in terms of accuracy is highlighted in **bold**, while the second-best configuration is indicated using *<u>italicized and underlined</u>* text.

**TABLE S8:**
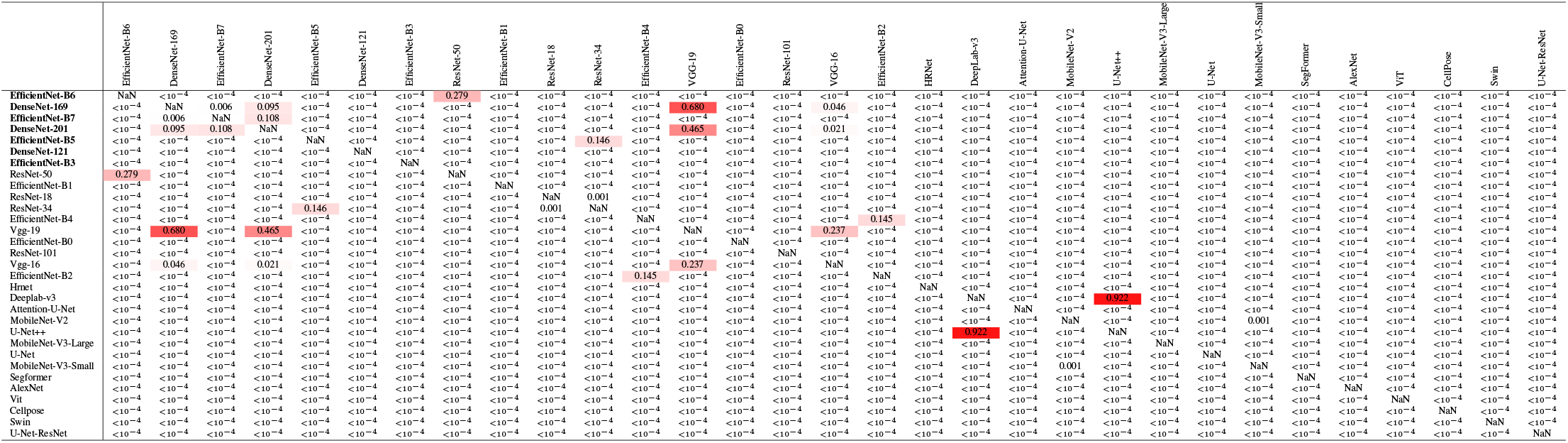
Pairwise Wilcoxon signed-rank test *p*-values computed from per-sample log-losses for all evaluated models. Values greater than 0.05 indicate no statistically significant difference in predictive confidence between the corresponding model pair.

## REFERENCES

[1] X. Xiong, L.-W. Zheng, Y. Ding, Y.-F. Chen, Y.-W. Cai, L.-P. Wang, L. Huang, C.-C. Liu, Z.-M. Shao, and K.-D. Yu, “Breast cancer: pathogenesis and treatments,” Signal transduction and targeted therapy, vol. 10, no. 1, p. 49, 2025.

[2] J. A. Aguirre-Ghiso, “Models, mechanisms and clinical evidence for cancer dormancy,” Nature Reviews Cancer, vol. 7, no. 11, pp. 834–846, 2007.

[3] M. A. Summers, M. M. McDonald, and P. I. Croucher, “Cancer cell dormancy in metastasis,” Cold Spring Harbor Perspectives in Medicine, vol. 10, no. 4, p. a037556, 2020.

[4] M. S. Sosa, P. Bragado, and J. A. Aguirre-Ghiso, “Mechanisms of disseminated cancer cell dormancy: an awakening field,” Nature Reviews Cancer, vol. 14, no. 9, pp. 611–622, 2014.

[5] R. N. Pedersen, B. Ö. Esen, L. Mellemkjær, P. Christiansen, B. Ejlertsen, T. L. Lash, M. Nørgaard, and D. Cronin-Fenton, “The incidence of breast cancer recurrence 10-32 years after primary diagnosis,” JNCI: Journal of the National Cancer Institute, vol. 114, no. 3, pp. 391–399, 2022.

[6] H. Hosseini, M. M. Obradović, M. Hoffmann, K. L. Harper, M. S. Sosa, M. Werner-Klein, L. K. Nanduri, C. Werno, C. Ehrl, M. Maneck et al., “Early dissemination seeds metastasis in breast cancer,” Nature, vol. 540, no. 7634, pp. 552–558, 2016.

[7] C. M. Koebel, W. Vermi, J. B. Swann, N. Zerafa, S. J. Rodig, L. J. Old, M. J. Smyth, and R. D. Schreiber, “Adaptive immunity maintains occult cancer in an equilibrium state,” Nature, vol. 450, no. 7171, pp. 903–907, 2007.

[8] I. Heidrich, B. Deitert, S. Werner, and K. Pantel, “Liquid biopsy for monitoring of tumor dormancy and early detection of disease recurrence in solid tumors,” Cancer and Metastasis Reviews, vol. 42, no. 1, pp. 161–182, 2023.

[9] S. Sauer, D. R. Reed, M. Ihnat, R. E. Hurst, D. Warshawsky, and D. Barkan, “Innovative approaches in the battle against cancer recurrence: novel strategies to combat dormant disseminated tumor cells,” Frontiers in oncology, vol. 11, p. 659963, 2021.

[10] D. Barkan, J. E. Green, and A. F. Chambers, “Extracellular matrix: a gatekeeper in the transition from dormancy to metastatic growth,” European journal of cancer, vol. 46, no. 7, pp. 1181–1188, 2010.

[11] D. Barkan, H. Kleinman, J. L. Simmons, H. Asmussen, A. K. Kamaraju, M. J. Hoenorhoff, Z.-y. Liu, S. V. Costes, E. H. Cho, S. Lockett et al., “Inhibition of metastatic outgrowth from single dormant tumor cells by targeting the cytoskeleton,” Cancer research, vol. 68, no. 15, pp. 6241–6250, 2008.

[12] D. Barkan and J. E. Green, “An in vitro system to study tumor dormancy and the switch to metastatic growth,” Journal of Visualized Experiments: JoVE, no. 54, p. 2914, 2011.

[13] N. Linde, G. Fluegen, and J. Aguirre-Ghiso, “The relationship between dormant cancer cells and their microenvironment,” Advances in cancer research, vol. 132, pp. 45–71, 2016.

[14] K. Kansal, S. Kumar, and K. Kansal, “Advances in deep learning techniques for breast cancer classification: A comprehensive review: K. kansal,” Archives of Computational Methods in Engineering, vol. 33, no. 1, pp. 187–222, 2026.

[15] K. He, X. Zhang, S. Ren, and J. Sun, “Deep residual learning for image recognition,” in Proceedings of the IEEE conference on computer vision and pattern recognition, 2016, pp. 770–778.

[16] M. Tan and Q. Le, “Efficientnet: Rethinking model scaling for convolutional neural networks,” in International conference on machine learning. PMLR, 2019, pp. 6105–6114.

[17] P. Rajpurkar, J. Irvin, K. Zhu, B. Yang, H. Mehta, T. Duan, D. Ding, A. Bagul, C. Langlotz, K. Shpanskaya et al., “Chexnet: Radiologist-level pneumonia detection on chest x-rays with deep learning,” arXiv preprint 1711.05225, 2017.

[18] S. Ben Baruch, N. Rotman-Nativ, A. Baram, H. Greenspan, and N. T. Shaked, “Cancer-cell deep-learning classification by integrating quantitative-phase spatial and temporal fluctuations,” Cells, vol. 10, no. 12, p. 3353, 2021.

[19] C. Piansaddhayanon, C. Koracharkornradt, N. Laosaengpha, Q. Tao, P. Ingrungruanglert, N. Israsena, E. Chuangsuwanich, and S. Sriswasdi, “Label-free tumor cells classification using deep learning and high-content imaging,” Scientific Data, vol. 10, no. 1, p. 570, 2023.

[20] O. Ronneberger, P. Fischer, and T. Brox, “U-net: Convolutional networks for biomedical image segmentation,” in International Conference on Medical image computing and computer-assisted intervention. Springer, 2015, pp. 234–241.

[21] L.-C. Chen, G. Papandreou, F. Schroff, and H. Adam, “Rethinking atrous convolution for semantic image segmentation,” arXiv preprint 1706.05587, 2017.

[22] K. Sun, Y. Zhao, B. Jiang, T. Cheng, B. Xiao, D. Liu, Y. Mu, X. Wang, W. Liu, and J. Wang, “High-resolution representations for labeling pixels and regions,” arXiv preprint 1904.04514, 2019.

[23] A. Dosovitskiy, L. Beyer, A. Kolesnikov, D. Weissenborn, X. Zhai, T. Unterthiner, M. Dehghani, M. Minderer, G. Heigold, S. Gelly et al., “An image is worth 16×16 words: Transformers for image recognition at scale,” arXiv preprint 2010.11929, 2020.

[24] Z. Liu, Y. Lin, Y. Cao, H. Hu, Y. Wei, Z. Zhang, S. Lin, and B. Guo, “Swin transformer: Hierarchical vision transformer using shifted windows,” in Proceedings of the IEEE/CVF international conference on computer vision, 2021, pp. 10012–10022.

[25] E. Xie, W. Wang, Z. Yu, A. Anandkumar, J. M. Alvarez, and P. Luo, “Segformer: Simple and efficient design for semantic segmentation with transformers,” Advances in neural information processing systems, vol. 34, pp. 12077–12090, 2021.

[26] A. Kirillov, E. Mintun, N. Ravi, H. Mao, C. Rolland, L. Gustafson, T. Xiao, S. Whitehead, A. C. Berg, W.-Y. Lo et al., “Segment anything,” in Proceedings of the IEEE/CVF international conference on computer vision, 2023, pp. 4015–4026.

[27] S. Woo, S. Debnath, R. Hu, X. Chen, Z. Liu, I. S. Kweon, and S. Xie, “Convnext v2: Co-designing and scaling convnets with masked autoencoders,” in Proceedings of the IEEE/CVF conference on computer vision and pattern recognition, 2023, pp. 16133–16142.

[28] C. Dong, Y. Liu, S. Chong, J. Zeng, Z. Bian, X. Chen, and S. Fan, “Deciphering dormant cells of lung adenocarcinoma: prognostic insights from o-glycosylation-related tumor dormancy genes using machine learning,” International Journal of Molecular Sciences, vol. 25, no. 17, p. 9502, 2024.

[29] L. A. Quayle, A. Spicer, P. D. Ottewell, and I. Holen, “Transcriptomic profiling reveals novel candidate genes and signalling programs in breast cancer quiescence and dormancy,” Cancers, vol. 13, no. 16, p. 3922, 2021.

[30] J. Xie, R. Liu, J. Luttrell IV, and C. Zhang, “Deep learning based analysis of histopathological images of breast cancer,” Frontiers in genetics, vol. 10, p. 80, 2019.

[31] A. Boukaache, B. N. Edinne, and D. Boudjehem, “Breast cancer image classification using convolutional neural networks (cnn) models,” International Journal of Informatics and Applied Mathematics, vol. 6, no. 2, pp. 20–34, 2024.

[32] M. Nasser and U. K. Yusof, “Deep learning based methods for breast cancer diagnosis: a systematic review and future direction,” Diagnostics, vol. 13, no. 1, p. 161, 2023.

[33] H. Li, V. Govindarajan, T. F. Ang, Z. A. Shaikh, A. Ksibi, Y.-L. Chen, C. S. Ku, M. C. Leong, F. H. Shabaruddin, W. Z. Wan Ishak et al., “Mspo: A machine learning hyperparameter optimization method for enhanced breast cancer image classification,” Digital Health, vol. 11, p. 20552076251361603, 2025.

[34] C. J. Ejiyi, D. Cai, D. L. Fiasam, B. Adjei-Arthur, S. Obiora, B. J. Ayekai, S. K. Asare, A. L. Jonathan, and Z. Qin, “Multi-modality medical image classification with resomergenet for cataract, lung cancer, and breast cancer diagnosis,” Computers in Biology and Medicine, vol. 187, p. 109791, 2025.

[35] D. Zhang, L. Dihge, P.-O. Bendahl, I. Arvidsson, M. Dustler, J. Ellbrant, K. Gulis, M. Hjärtström, M. Ohlsson, C. Rejmer et al., “Deep learning on routine full-breast mammograms enhances lymph node metastasis prediction in early breast cancer,” npj Digital Medicine, vol. 8, no. 1, p. 425, 2025.

[36] Q. Zhang, H. Gao, W. Li, Z. Xu, T. Ouyang, and Z. Gu, “Enhanced nuclear information fusion and visual transformer for pathological breast cancer image classification,” Scientific Reports, vol. 15, no. 1, p. 19490, 2025.

[37] S. Malik, S. G. K. Patro, A. K. J. Al-Nussairi, C. Mahanty, M. Ghouse, A. Thiyagarajan, A. A. Hadi, A. Khan, M. Mittal, and A. Zewude, “A unified multi modal transformer framework for breast cancer recurrence prediction and survival analysis,” Scientific Reports, 2026.

[38] S. Elumalai, S. Rajendran, and M. Khalid, “Breast cancer classification based on microcalcifications using dual branch vision transformer fusion,” Scientific Reports, 2025.

[39] M. M. Ali, A. Shabbir, A. Akbar, A. Angelopoulou, M. Aftab, Q. Zia, C. Wang, and W. Zhenfei, “Ctnet: multi-modal channel attention transformer network for breast cancer image classification,” Biomedical Signal Processing and Control, vol. 113, p. 109108, 2026.

[40] I. Abdelsabour, A. Elgarayhi, M. Sallah, and M. Elmogy, “Different birads breast cancer diagnosis using mobilenetv1 and vision transformer based on explainable artificial intelligence (xai),” Scientific Reports, 2026.

[41] A. Brahmareddy and M. P. Selvan, “Transbreastnet a cnn transformer hybrid deep learning framework for breast cancer subtype classification and temporal lesion progression analysis,” Scientific Reports, vol. 15, no. 1, p. 35106, 2025.

[42] I. Jahan, M. E. Chowdhury, S. Vranic, R. M. Al Saady, S. Kabir, Z. H. Pranto, S. J. Mim, and S. F. Nobi, “Deep learning and vision transformers-based framework for breast cancer and subtype identification,” Neural Computing and Applications, vol. 37, no. 16, pp. 9311–9330, 2025.

[43] A. Vaswani, N. Shazeer, N. Parmar, J. Uszkoreit, L. Jones, A. N. Gomez, Ł. Kaiser, and I. Polosukhin, “Attention is all you need,” in Advances in neural information processing systems, 2017, pp. 5998–6008.

[44] G. Huang, Z. Liu, L. Van Der Maaten, and K. Q. Weinberger, “Densely connected convolutional networks,” in Proceedings of the IEEE conference on computer vision and pattern recognition, 2017, pp. 4700–4708.

[45] M. Sandler, A. Howard, M. Zhu, A. Zhmoginov, and L.-C. Chen, “Mobilenetv2: Inverted residuals and linear bottlenecks,” in Proceedings of the IEEE conference on computer vision and pattern recognition, 2018, pp. 4510–4520.

[46] A. Howard, M. Sandler, G. Chu, L.-C. Chen, B. Chen, M. Tan, W. Wang, Y. Zhu, R. Pang, V. Vasudevan et al., “Searching for mobilenetv3,” in Proceedings of the IEEE/CVF international conference on computer vision, 2019, pp. 1314–1324.

[47] Z. Zhou, M. M. Rahman Siddiquee, N. Tajbakhsh, and J. Liang, “Unet++: A nested u-net architecture for medical image segmentation,” in International workshop on deep learning in medical image analysis. Springer, 2018, pp. 3–11.

[48] O. Oktay, J. Schlemper, L. L. Folgoc, M. Lee, M. Heinrich, K. Misawa, K. Mori, S. McDonagh, N. Y. Hammerla, B. Kainz et al., “Attention u-net: Learning where to look for the pancreas,” arXiv preprint 1804.03999, 2018.

[49] C. Stringer, T. Wang, M. Michaelos, and M. Pachitariu, “Cellpose: a generalist algorithm for cellular segmentation,” Nature methods, vol. 18, no. 1, pp. 100–106, 2021.

